# HCN channel-mediated neuromodulation can control action potential velocity and fidelity in central axons

**DOI:** 10.1101/643197

**Authors:** Niklas Byczkowicz, Abdelmoneim Eshra, Jacqueline Montanaro, Andrea Trevisiol, Johannes Hirrlinger, Maarten H. P. Kole, Ryuichi Shigemoto, Stefan Hallermann

## Abstract

Hyperpolarization-activated cyclic-nucleotide-gated (HCN) channels control electrical rhythmicity and excitability in the heart and brain, but the function of HCN channels at subcellular level in axons remains poorly understood. Here, we show that the action potential conduction velocity in both myelinated and unmyelinated central axons can bidirectionally be modulated by HCN channel blockers, cyclic adenosine monophosphate (cAMP), and neuromodulators. Recordings from mice cerebellar mossy fiber boutons show that HCN channels ensure reliable high-frequency firing and are strongly modulated by cAMP (EC_50_ 40 µM; estimated endogenous cAMP concentration 13 µM). In accord, immunogold-electron microscopy revealed HCN2 as the dominating subunit in cerebellar mossy fibers. Computational modeling indicated that HCN2 channels control conduction velocity primarily via altering the resting membrane potential and was associated with significant metabolic costs. These results suggest that the cAMP-HCN pathway provides neuromodulators an opportunity to finely tune energy consumption and temporal delays across axons in the brain.

## Introduction

HCN channels are expressed in the heart and nervous system and comprise four members (HCN1–HCN4) differing in their kinetics, voltage-dependence and degree of sensitivity to cyclic nucleotides such as cAMP (Biel et al., 2009; Robinson and Siegelbaum, 2003). Membrane hyperpolarization activates HCN channels and causes a depolarizing mixed sodium/potassium (Na^+^/K^+^) current. In the heart, the current through HCN channels (*I*_f_) mediates the acceleratory effect of adrenaline on heart rate by direct binding of cAMP (DiFrancesco, 2006). In neurons, the current through HCN channels (*I*_h_) controls a wide array of functions, such as rhythmic activity (Pape and McCormick, 1989) and excitability (Tang and Trussell, 2015). In addition to the somatic impact, HCN channels are expressed throughout various subcellular compartments of neurons (Nusser, 2012). For example, patch-clamp recordings from dendrites in pyramidal neurons have revealed particularly high densities of HCN channels which acts to control the local resting potential and leak conductance, thereby playing important roles in regulating synaptic integration (George et al., 2009; Harnett et al., 2015; Kole et al., 2006; Magee, 1999; Williams and Stuart, 2000).

In contrast, the expression and role of *I*_h_ in the axon is less studied and may vary with species. *I*_h_ seems to critically control the strength of synaptic transmission in crayfish and *Drosophila* neuromuscular junction (Beaumont and Zucker, 2000; Cheung et al., 2006). However, presynaptic recordings from the vertebrate calyx of Held in the auditory brainstem found *I*_h_ to marginally affect neurotransmitter release (Cuttle et al., 2001), but to exert a strong influence on the resting membrane potential (Cuttle et al., 2001; Kim and von Gersdorff, 2012) and vesicular neurotransmitter uptake (Huang and Trussell, 2014). At synaptic terminals of pyramidal neurons in the cortex of mice, HCN channels inhibit glutamate release by suppressing the activity of T-type Ca^2+^ channels (Huang et al., 2011).

Besides a potential impact on neurotransmitter release, axonal *I*_h_ could play a role in the propagation of action potentials. Indeed, in axons of the stomatogastric nervous system of lobsters (Marder and Bucher, 2001) it was shown that the action potential conduction was affected by dopamine via axonal HCN channels (Ballo et al., 2010; Ballo et al., 2012). In vertebrates, studies on action potential propagation by Waxman and coworkers indicated that *I*_h_ counteracts the hyperpolarization of the membrane potential during periods of high-frequency firing (Baker et al., 1987; Birch et al., 1991; Waxman et al., 1995) and that it participates in ionic homeostasis at the node of Ranvier (Waxman and Ritchie, 1993). More recent investigations found *I*_h_ to be crucial for the emergence of persistent action potential firing in axons of parvalbumin-positive interneurons (Elgueta et al., 2015), but *I*_h_ seems to have an opposing effect on the excitability at the axon initial segment, where its activation reduces the probability of action potential initiation (Ko et al., 2016). Finally, there is evidence from extracellular recordings that blocking *I*_h_ decreases the action potential conduction velocity in unmyelinated central (Baginskas et al., 2009; Soleng et al., 2003) and peripheral axons of vertebrates (Grafe et al., 1997). However, neuromodulation of conduction velocity and the underlying cellular membrane mechanisms are not known in vertebrate axons.

Here, we demonstrate a decrease or increase in conduction velocity in central axons through the application of HCN blockers or neuromodulators. To gain mechanistic insights into the modulation of conduction velocity by HCN channels, we performed recordings from *en passant* cerebellar mossy fiber boutons (cMFB; Ritzau-Jost et al., 2014; Delvendahl et al., 2015). We found that HCN channels in cMFBs mainly consist of the HCN2 subunit, are ∼7% activated at resting membrane potential, ensure high-frequency firing, and control the passive membrane properties. Whole-cell and perforated patch clamp recordings from cMFBs demonstrated a strong dependence of HCN channels on intracellular cAMP concentration with an EC_50_ of 40 µM and a high endogenous cAMP concentration of 13 µM. Computational modelling indicated that the resting membrane potential controls conduction velocity and that the activity of HCN channel is metabolically expensive. These data reveal a mechanism shared among different types of axons to bidirectionally modulate conduction velocity in the central nervous system.

## Results

### Bidirectional modulation of conduction velocity

To investigate whether HCNs affect conduction velocity, we recorded compound action potentials in three different types of axons (Fig. 1): Application of the specific HCN channel blocker ZD7288 (30 µM) decreased the conduction velocity by 8.0 ± 2.8% in myelinated cerebellar mossy fibers (n = 14), by 9.2 ± 0.9% in non-myelinated cerebellar parallel fibers (n = 15), and by 4.0 ± 0.8% in optical nerves (n = 4; see Fig. 1 and legend for statistical testing). Since some studies implied that ZD7288 might have some unspecific side effects, such as blocking voltage-dependent Na^+^ channels (Chevaleyre and Castillo, 2002; Wu et al., 2012), we recorded Na^+^ currents from 53 cMFBs and found no change in amplitude or kinetics of voltage-dependent Na^+^ currents after ZD7288 application (Supplemental Fig. 1), suggesting that under our conditions and at a concentration of 30 µM, ZD7288 did not affect the Na^+^ currents. Because of the modulation of HCN channels by intracellular cAMP, we measured conduction velocity during application of 8-bromoadenosine 3′,5′-cyclic monophosphate (8-Br-cAMP; 500 µM), a membrane-permeable cAMP-analogue. The conduction velocity increased by 5.9 ± 2.8% in cerebellar mossy fibers (n = 17), by 3.7 ± 1.4% in parallel fibers (n = 10), and by 4.6 ± 0.6% in optic nerves (n = 5; see Fig. 1 and legend for statistical testing). These results indicate that HCN channels control the conduction velocity both in myelinated and non-myelinated central axons.

**Figure 1.**
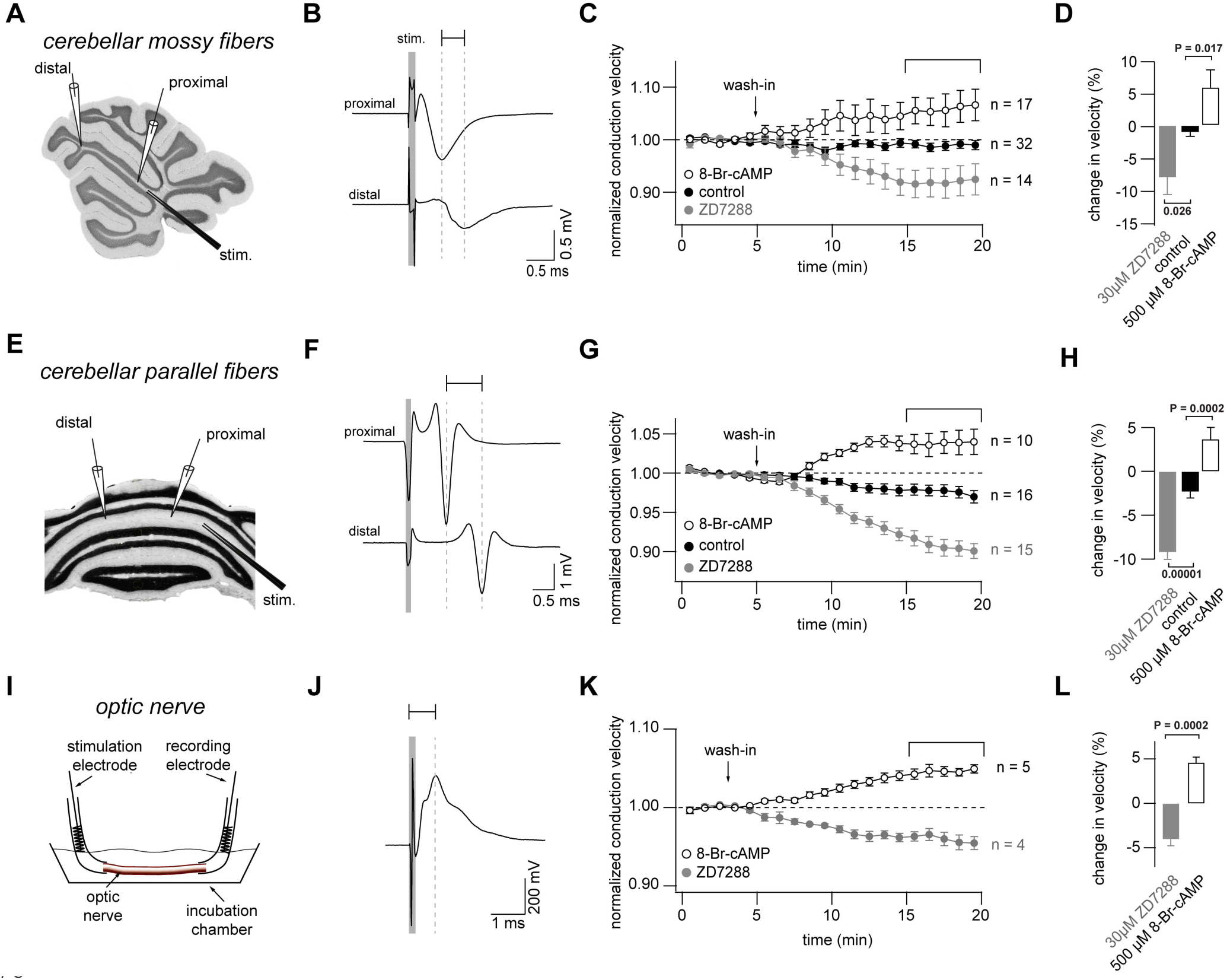
Bidirectional modulation of conduction velocity. (A) Recording configuration of conduction velocity in mossy fibers using a bipolar tungsten stimulation electrode (stim.) and two glass electrodes filled with 1M NaCl solution. (B) Example of compound action potentials recorded with two electrodes positioned with different distance in relation to the stimulation electrode. Stimulation was 100 µs as indicated by the grey bar. Each trace is an average of 50 individual compound action potentials recorded at 1Hz. (C) Average normalized mossy fiber conduction velocity, during bath application ZD7288 (30 µM) or 8-Br-cAMP (500 µM) at t = 5 min. (D) Average relative change in mossy fiber conduction velocity after application of ZD7288 or 8-Br-cAMP measured 10 to 15 minutes after wash-in (bracket in C). P_ANOVA_ = 0.00015. P_Kruskal-Wallis_ = 0.00044. The individual P values of the Dunnett test for multiple comparisons with control are indicated. (E) Schematic illustration of the experimental configuration used to record from cerebellar parallel fibers. (F) Examples of compound action potentials recorded from parallel fibers, as in panel B. (G) Normalized conduction velocity in parallel fibers, as in panel C. (H) Average relative changes in parallel fiber, as in panel D. P_ANOVA_ = 10^−9^. P_Kruskal-Wallis_ = 10^−8^. The individual P values of the Dunnett test for multiple comparisons with control are indicated. (I) Schematic of the experimental configuration used to record from optic nerve (J) Examples of compound action potentials recorded from optic nerve, as in panel B. (K) Normalized conduction velocity in optic nerve, as in panel C. (L) Average relative changes in optic nerve, as in panel D. P_T-Test_ = 0.0002. P_Wilcoxon-Mann-Whitney-Test_ = 0.004.

### Neuromodulators differentially regulate conduction velocity

To investigate a modulation of conduction velocity by physiological neuromodulators, we focused on the cerebellar parallel fibers, where the velocity could be most accurately measured, and then applied several modulators known to act via cAMP-dependent pathways (Fig. 2). Wash-in of 200 µM norepinephrine (NE) resulted in a relatively fast increase in conduction velocity (1.9 ± 0.8%; n = 6; see Fig. 2B and legend for statistical testing), consistent with the existence of β-adrenergic receptors in the cerebellar cortex (Nicholas et al., 1993), which increase the cAMP concentration via G_s_-proteins. On the other hand, the application of either 200 µM serotonin (–3.5 ± 0.5%; n = 11), 200 µM dopamine (–5.0 ± 0.7%; n = 13) or 200 µM adenosine (–7.2 ± 0.6%; n = 5) resulted in a continuous decrease of the conduction velocity (Fig. 2B and C; see legend for statistical testing), consistent with the existence of G_i_-coupled receptors for serotonin, dopamine, and adenosine in the molecular layer of the cerebellum (Geurts et al., 2002; Schweighofer et al., 2004), which decrease the cAMP concentration. Although we used rather high concentrations of the agonists and off-target effects cannot be excluded (e.g., NE activating dopamine receptors; Sánchez-Soto et al., 2016), these data nevertheless indicate that physiological neuromodulators can both increase or decrease action potential conduction velocity, depending on the type of neuromodulator and receptor.

**Figure 2.**
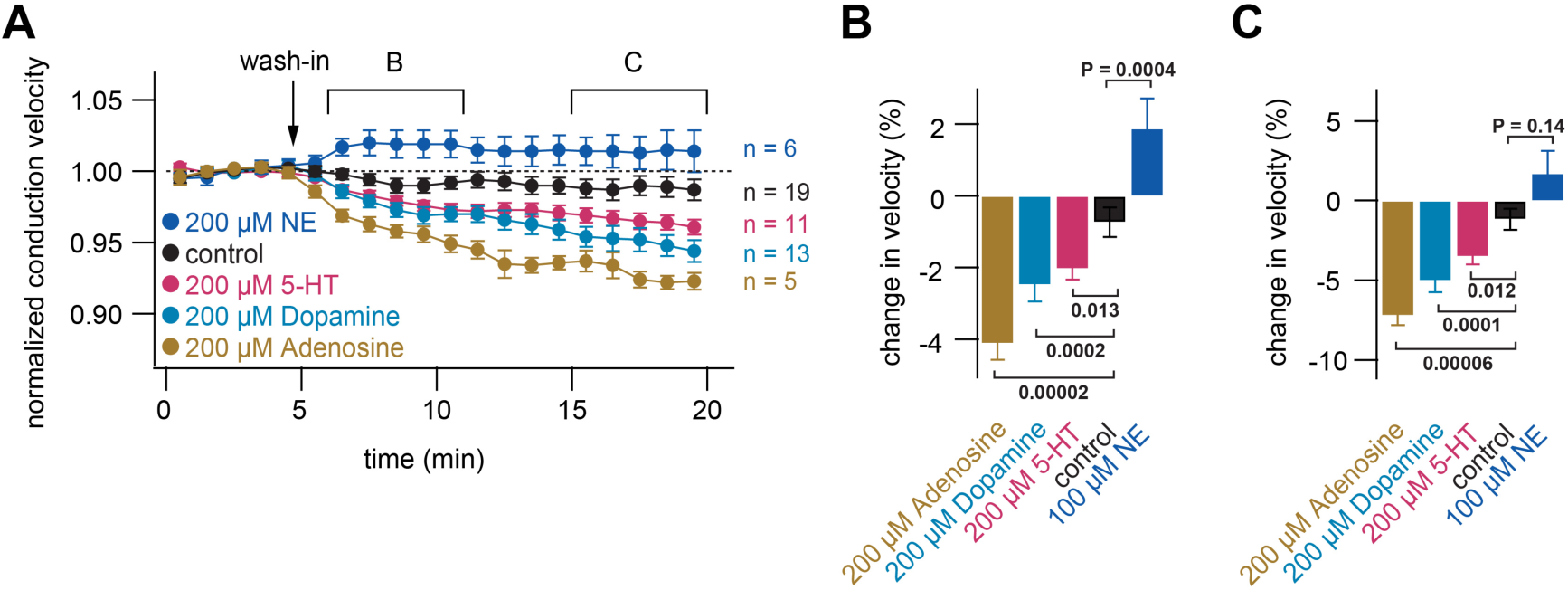
Neuromodulators differentially regulate conduction velocity. (A) Average normalized conduction velocity in cerebellar parallel fibers during wash-in at t = 5 min of various neuromodulators known to act via cAMP-dependent pathways. (B) Average relative change in conduction velocity after application of the neuromodulators measured from 1 to 6 minutes after wash-in (bracket marked B in panel A). P_ANOVA_ = 9*10^−10^. P_Kruskal-Wallis_ = 3*10^−8^. The individual P values of the Dunnett test for multiple comparisons with control are indicated. (C) Average relative change in conduction velocity after application of the neuromodulators measured 10 to 15 minutes after wash-in (bracket marked C in panel A). P_ANOVA_ = 3*10^−7^. P_Kruskal-Wallis_ = 3*10^−7^. The individual P values of the Dunnett test for multiple comparisons with control are indicated.

### Neuromodulation of conduction velocity is mediated by HCN channels

In addition to HCN channels, some voltage gated Na^+^, K^+^, and Ca^2+^ channels can be modulated via the intracellular cAMP-pathway (Burke et al., 2018; Yang et al., 2013; Yin et al., 2017). To address the contribution of other channels on the neuromodulation of the conduction velocity, we performed a set of experiments, in which HCN channels were first blocked by 30 µM ZD7288 and subsequently three modulatory substances that significantly increased or decreased conduction velocity in prior experiments were applied. With ZD7288 continuously being present in the recording solution, conduction velocity of parallel fibers slightly decreased over the course of 20 minutes (Fig. 3A). Adding 8Br-cAMP (500 µM), Adenosine (200 µM) or NE (100 µM) at t = 5 min (i.e. 25 minutes after application of ZD7288) did not change the conduction velocity significantly compared with control (only ZD7288). The average conduction velocity between t = 15 and 20 min was decreased by –3.3 ± 2.4% for cAMP (n = 9), –4.6 ± 1.6% for Adenosine (n = 9) and –3.7 ± 1.2% for NE (n = 7), compared to the average velocity between t = 0 and 5 min baseline recording. This was not significantly different from the decrease in the sole presence of ZD7288 (–3.3 ± 1.4%; n = 7, see Fig. 3B and legend for statistical testing), indicating that the previously shown effects of cAMP and neuromodulators on conduction velocity are mainly mediated by HCN channels.

**Figure 3.**
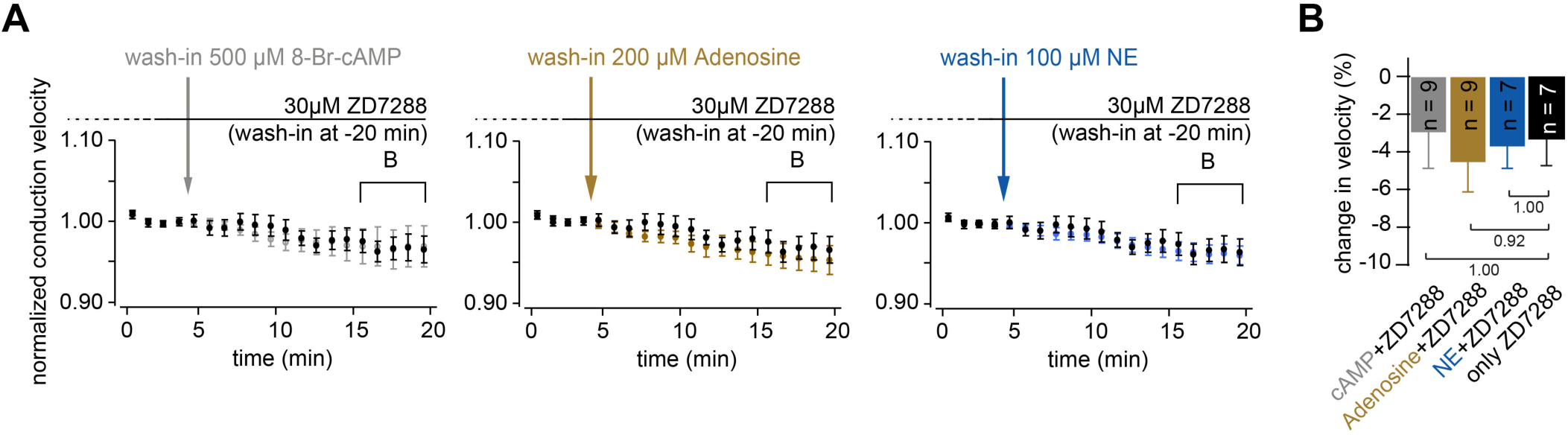
Neuromodulation of conduction velocity is mediated by HCN channels. (A) Average normalized conduction velocity in parallel fibers. Wash-in of 30 µM ZD7288 20 minutes before the start of the recording. The substance remained in the solution during recording to ensure continuous block HCN channels. At t = 5 min, 8-Br-cAMP, Adenosine or NE was added to the solution. (B) Average relative change in conduction velocity after application of the neuromodulators measured 10 to 15 minutes after wash-in (bracket marked B in panel A). P_ANOVA_ = 0.91. P_Kruskal-Wallis_ = 0.77. The individual P values of the Dunnett test for multiple comparisons with control are indicated.

### Cerebellar mossy fiber terminals have a prominent voltage sag

To investigate the membrane and signaling mechanisms underlying the bidirectional control of conduction velocity, we focused on cerebellar mossy fibers, which allow the whole-cell recording configuration and a direct access to the cytoplasmic compartment (Fig. 4A). Recordings from *en passant* cMFBs are well suited to investigate the ionic basis of conduction velocity in the adjacent axonal compartments, because of a long membrane length constant and slow HCN channel gating. Injection of depolarizing currents during current-clamp recordings evoked a single action potential and injection of hyperpolarizing currents generated a substantial ‘sag’ at cMFBs (Fig. 4B; Rancz et al., 2007; Ritzau-Jost et al., 2014), i.e. a delayed depolarization towards the resting potential, which is a hallmark of the presence of *I*_h_ (Biel et al., 2009; Robinson and Siegelbaum, 2003). At a potential of on average –150 mV, the sag ratio, calculated from the peak and steady state amplitude as indicated in Fig. 4C (George et al., 2009), was 0.497 ± 0.030 (n = 12).

**Figure 4.**
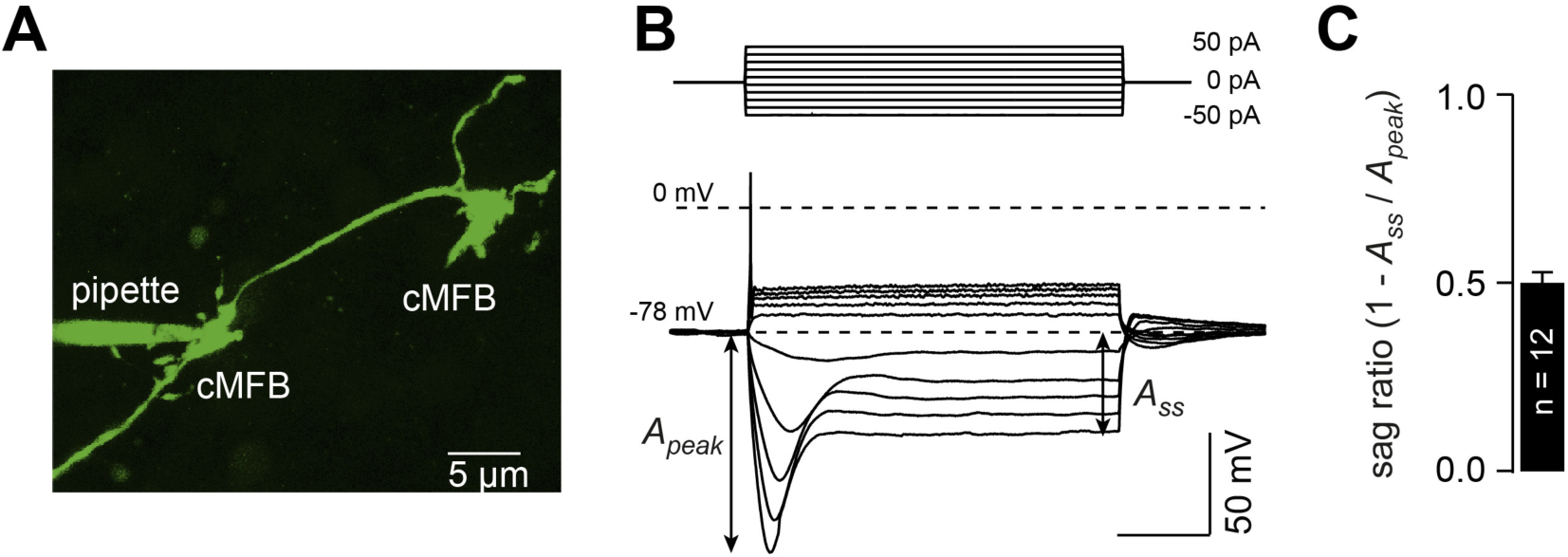
Cerebellar mossy fiber terminals have a prominent voltage sag. (A) Two-photon microscopic image of a whole-cell patch-clamp recording from a cMFB (green) filled with the fluorescence dye Atto 488 in an acute cerebellar brain slice of an adult 39-days old C57/Bl6 mouse (maximal projection of stack of images). (B) Characteristic response of a cMFB to current injection: Depolarizing pulses evoked a single action potential while hyperpolarizing pulses evoked a strong hyperpolarization with a sag. (C) Average sag ratio of 12 cMFB recordings.

### HCN channels support high frequency action potential firing

Using direct presynaptic recordings from cMFBs, we first aimed to investigate the impact of HCN channels on action potential firing. To this end, we analyzed action potentials elicited by current injections into the cMFBs (data not shown) as well as traveling action potentials elicited by axonal stimulation with a second pipette (Fig. 5A). In both cases, the amplitude and half-duration of action potentials elicited at 1 Hz were not significantly affected by application of 30 µM ZD7288 (data not shown and Fig. 5B and C, respectively), indicating that HCN channels do not alter the active membrane properties profoundly.

**Figure 5.**
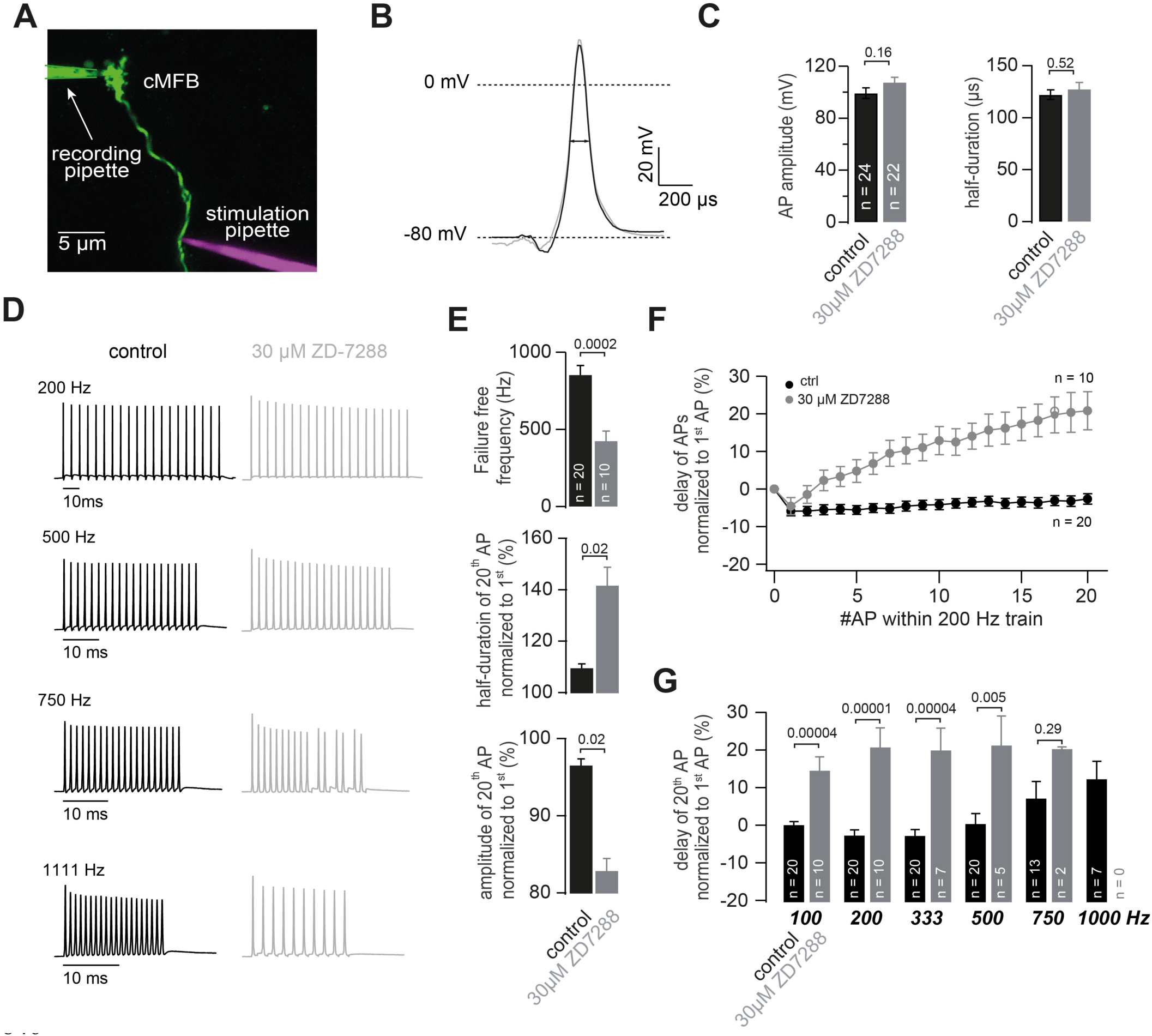
HCN channels support high frequency action potential firing. (A) Two-photon microscopic image of a whole-cell patch-clamp recording from a cMFB (green) filled with the fluorescence dye Atto 488 in an acute cerebellar brain slice of an adult 43-days old mouse (maximal projection of stack of images). Targeted axonal stimulation was performed by adding a red dye Atto 594 to the solution of the stimulation pipette. (B) Grand average of action potentials evoked at 1 Hz under control conditions (black) and in the presence of ZD7288 (grey). (C) Average action potential amplitude (measured from resting to peak) and half-duration P_T-Test_ = 0.16 and 0.51, for amplitude and resting respectively. (D) ZD7288 effects the ability of cerebellar mossy fibers to fire high frequency action potentials: Examples of two different cMFBs stimulated at frequencies between 200 Hz and 1111 Hz under control conditions (*left, black*) or in the presence of 30 µM ZD7288 (*right, grey*). (E) cMFBs treated with ZD7288 showed an on average lower maximal failure-free firing frequency. In addition, amplitude reduction and action potential broadening during 200-Hz trains of action potentials in ZD7288 treated axons were more pronounced than under control conditions. (F) Average delay between the peak of the action potentials (AP) and the stimulation during 200 Hz trains of 20 action potentials normalized to the delay of the first action potential for control conditions (*black*) and in the presence of 30 µM ZD7288 (*grey*). (G) Average delay of the 20^th^ normalized to the delay of the 1^st^ action potential during trains of 20 action potentials at frequencies ranging from 100 to 1000 Hz. The provided P-values based on simple t-test are mostly much smaller than the Bonferroni-corrected significance left of 0.05/6 = 0.008, indicating a highly significant slowing of the conduction velocity during high-frequency trains.

However, cerebellar mossy fibers can conduct trains of action potentials at frequencies exceeding 1 kHz (Ritzau-Jost et al., 2014), making them an ideal target to study the impact of axonal HCNs on the propagation of high-frequency action potentials as well. Blocking HCN channels significantly impaired the ability of mossy fibers to fire at high frequencies (20 action potentials at 200 – 1666 Hz). In the examples illustrated in Fig. 5D, the failure-free trains of action potentials could be elicited at up to 1.1 kHz under control conditions and up to 500 Hz, when ZD7288 was present in the extracellular solution. The average failure-free frequency reduced from 854 ± 60 in control to 426 ± 63 Hz in the presence of ZD7288 (n = 20 and 10, respectively; P_T-Test_ = 0.0002; Fig. 5E). Action potential broadening and amplitude reduction was more pronounced in the presence of ZD7288. For example, during trains of action potentials at 200 Hz, the half-duration of the 20^th^ action potential was 109.6 ± 1.5 and 141.7 ± 7.0% of the half-duration of the 1^st^ action potential for control and ZD7288, respectively (n = 20 and 10; P_T-Test_ = 0.02; Fig. 5E). The amplitude of the 20^th^ action potential was 96.5 ± 0.8 and 82.9 ± 1.6% of the 1^st^ action potential for control and ZD7288, respectively (n = 20 and 10; P_T-Test_ = 0.02; Fig. 5E). Furthermore, the delay during trains of action potentials at 200 Hz increased by ∼20% in the presence of ZD7288 but decreased by ∼5% in control recordings (Fig. 5F), indicating an acceleration and a slowing of conduction velocity during high-frequency firing for control and ZD7288, respectively. The difference in delay of the 20^th^ action potential was maximal at intermediate frequencies (200 and 333 Hz; Fig. 5G). These experiments show, that HCNs, despite their slow kinetics, ensure reliable high-frequency firing.

### The passive membrane properties of cMFBs are HCN- and cAMP-dependent

To better understand how *I*_h_ impacts action potential firing, we next investigated the passive membrane properties of cMFBs by recording the voltage response elicited by small hyperpolarizing current injections (–10 pA for 300 ms) in the absence and presence of 30 µM ZD7288 (Fig. 6A and B). ZD7288 caused a hyperpolarization of the resting membrane potential by, on average, 5.4 mV (–80.0 ± 0.6 mV and –85.4 ± 1.4 mV for control and ZD7288, n = 94 and 35, respectively), a doubling of the apparent input resistance calculated from the steady state voltage at the end of the current step (794 ± 48 MΩ and 1681 ± 185 MΩ), as well as a doubling of the apparent membrane time constant, as determined by a mono-exponential fit to the initial decay of the membrane potential (14.4 ± 0.8 ms and 35.0 ± 2.5 ms, respectively; see legend of Fig. 6B for statistical testing). To analyze the cAMP-dependence of the conduction velocity (cf. Fig. 1), we determined the cAMP-dependence of the passive membrane properties of cMFBs. Adding cAMP in various concentrations to the intracellular solution depolarized the membrane potential and decreased both the input resistance and the apparent membrane time constant in a concentration-dependent manner, which shows an opposite effect compared to the application of ZD7288 (Fig. 6B). These data suggest that HCN channels in cerebellar mossy fibers determine the passive membrane properties as a function of the intracellular cAMP concentration.

**Figure 6.**
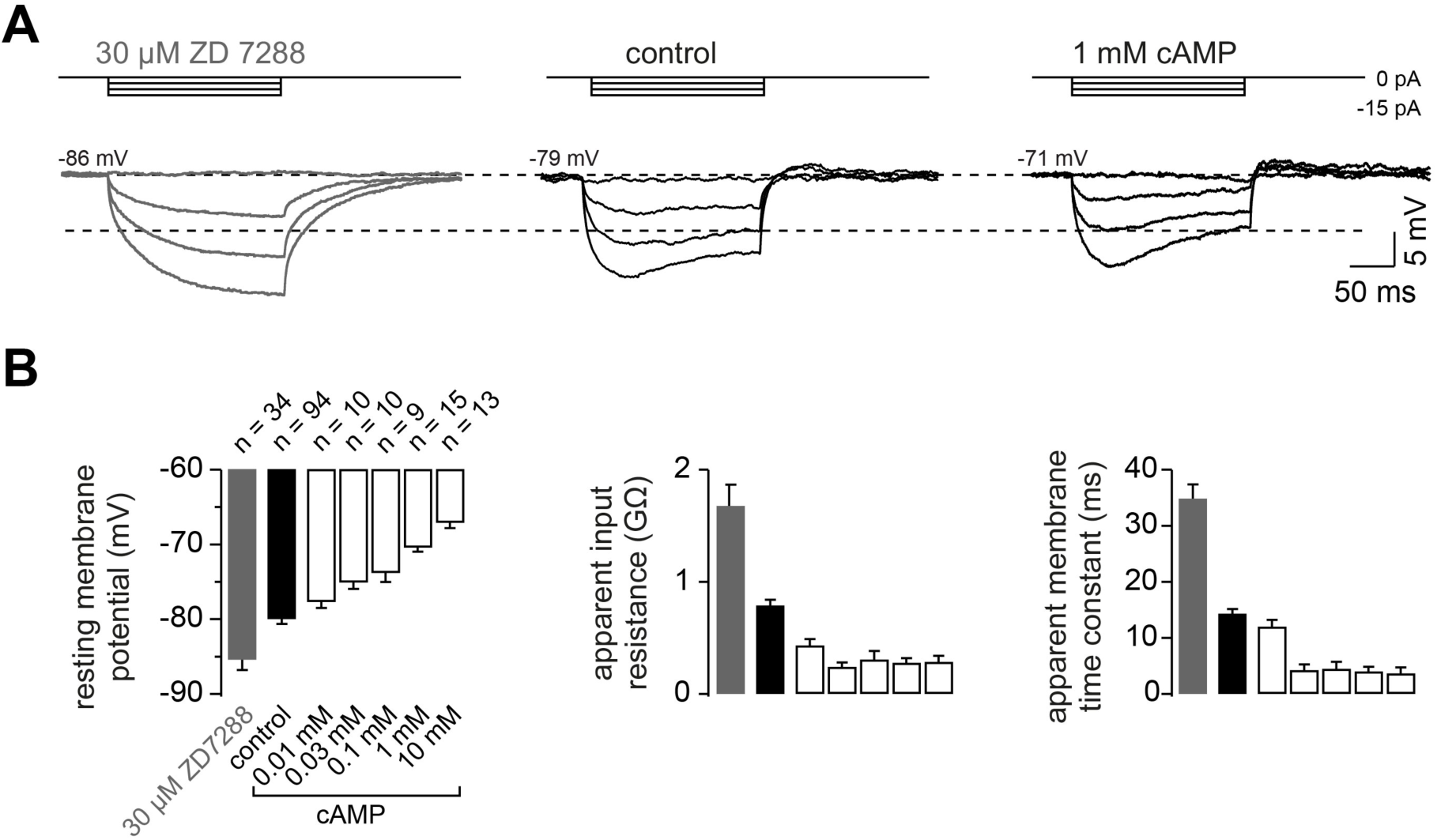
The passive membrane properties of cMFBs are HCN and cAMP-dependent. (A) Example voltage response of cMFBs to small hyperpolarizing current steps. Application of 30 µM ZD7288 eliminated the *I*_h_-mediated voltage sag (left). Adding 1 mM cAMP to the intracellular path-clamp solution (right) reduced the input resistance as seen by the reduced steady-state voltage response (dashed lines). (B) Average resting membrane potential (left), apparent input resistance (middle), and apparent membrane time constant (right) upon application of 30µM ZD7288 or different concentrations of cAMP. For all three parameters, P_ANOVA_ and P_Kruskal-Wallis_ are < 10^−10^ and the Dunnett test for multiple comparisons with control indicates, e.g., P < 0.0001 for control vs. ZD and P < 0.001 for control vs. 1 mM cAMP.

### HCN2 is uniformly distributed in mossy fiber axons and boutons

Of the four HCN subunits (HCN1–HCN4), the subunits HCN1 and HCN2 are predominantly expressed in the cerebellar cortex (Notomi and Shigemoto, 2004; Santoro et al., 2000). Previous studies in the cortex, hippocampus, and auditory brainstem primarily detected HCN1 in axons (Elgueta et al., 2015; Huang et al., 2011; Ko et al., 2016), but HCN2 was found to be more sensitive to cAMP in comparison to HCN1 (Wang et al., 2001; Zagotta et al., 2003). To understand the pronounced cAMP dependence of conduction velocity (cf. Fig. 1) and passive membrane properties (cf. Fig. 6) at the molecular level, we investigated the identity and distribution of HCN channels using pre-embedding immunogold labeling for HCN1 and 2 in cMFBs and adjacent axons. At the electron microscopic level, we found only background immunoreactivity for HCN1 (data not shown) but significant labeling for HCN2 (Fig. 7A). HCN2 immunogold particles were diffusely distributed along the plasma membrane of cMFBs, with similar labeling density in the adjacent mossy fiber axon (Fig. 7B), which could be traced back up to 3.5 µm from cMFBs. In addition, we created a 3D reconstruction of a cMFB (Fig. 7C and Supplemental Video), including gold particles for HCN2 and identified synaptic connections. While synapses onto granule cell dendrites were observed within invaginated parts of the bouton, HCN2 was uniformly distributed without apparent spatial relations within those synapses. The density of immunogold particles for HCN2 in this reconstructed bouton was 17.1 particles/µm^2^ (in total 1260 particles per 73.65 µm^2^). The mean density of immunogold particles for HCN2 was 22.7 ± 2.4 per μm^2^ (n = 6 cMFBs from 2 mice). These data indicate that HCN2 is the dominant subunit mediating *I*_h_ in cMFBs, consistent with its pronounced cAMP-dependence.

**Figure 7.**
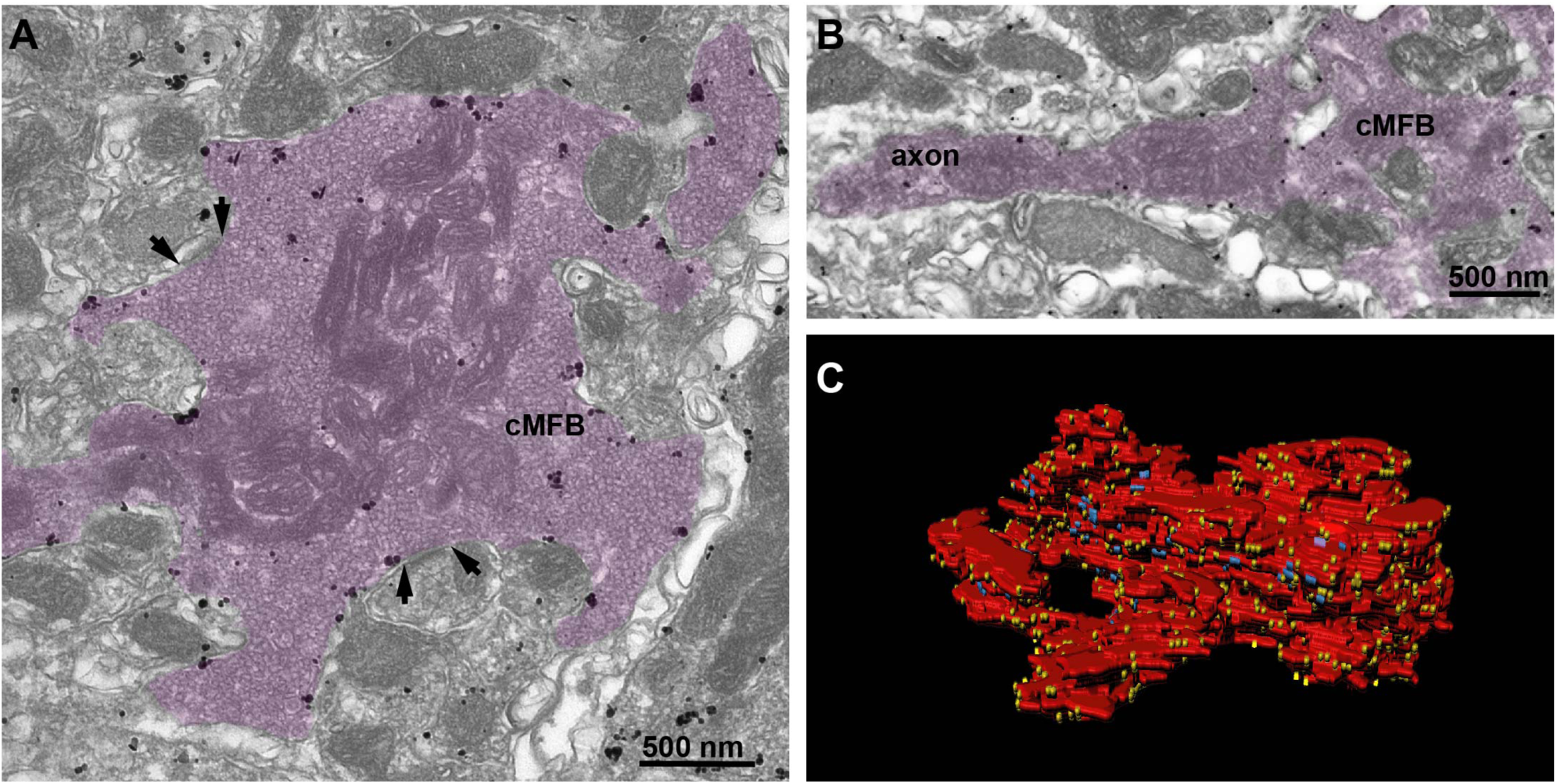
HCN2 is uniformly distributed in mossy fiber axons and boutons. (A) Electron microscopic image showing a cMFB (magenta) labeled for HCN2. Many particles are diffusely distributed along the plasma membrane of the cMFB, some of them being clustered. Arrows mark synapses between the cMFB and dendrites of adjacent GCs. (B) Another cMFB, showing similar labeling density for HCN2 in a proximal part of the mossy fiber axon. (C) Reconstructed cMFB (red) with identified synapses (blue) and HCN2 labeled with gold particles (yellow).

### HCN channels in cMFB are strongly modulated by cAMP

To better understand the function of axonal HCN2 channels and their modulation by intracellular cAMP, we performed voltage-clamp recordings from cMFBs combined with different cAMP concentrations within the intracellular patch solution. Hyperpolarizing voltage steps evoked a slowly activating, non-inactivating inward current, which was inhibited by ZD7288 (Fig. 8A). Using tail currents of ZD7288-sensitive currents evoked by voltage steps between –80 mV and –150 mV from a holding potential of –70 mV, we calculated the activation curve of *I*_h_ with a mean V_½_ of –103.3 ± 0.8 mV (Fig. 8B; n = 36 V_½_-values, each from a different cMFB). Based on the average resting membrane potential of cMFBs, this means about 7% of the overall HCN2-mediated current is active at rest.

**Figure 8.**
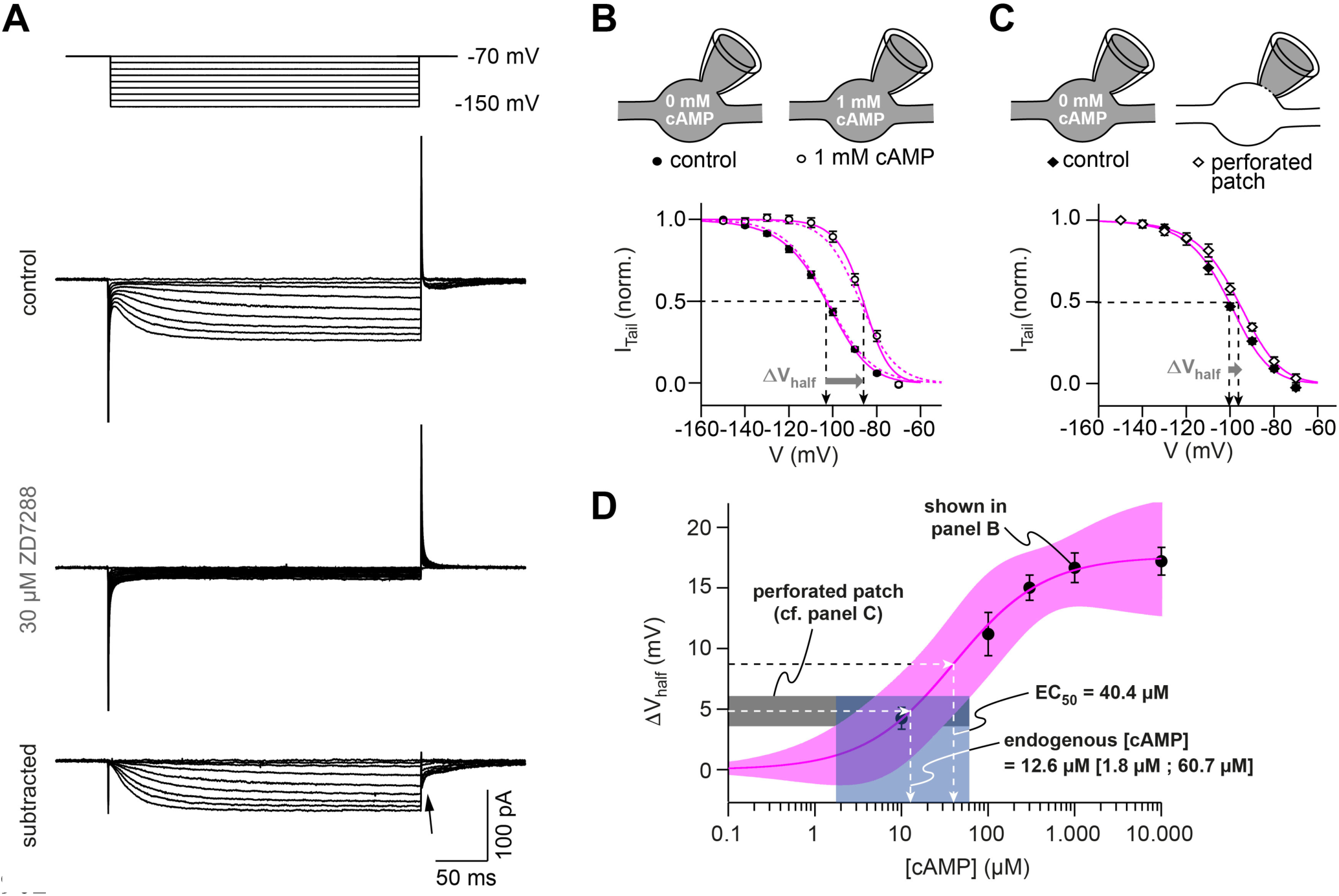
HCN channels in cMFB are strongly modulated by cAMP. (A) Example currents elicited hyperpolarizing voltage steps (–70 mV and then stepped to conditioning pulses between –80 mV and –150 mV). *Top*, the control current, *middle,* remaining transients in the presence of 30 µM ZD7288 and, *bottom*, the subtracted currents. The ZD7288-sensitive current is slowly activating, non-inactivating and shows inward tail currents (arrow). (B) Activation curve of *I*_h_ determined as the normalized tail current of ZD7288 sensitive currents obtained after the end of the conditioning voltage pulse (*arrow in* plotted versus corresponding voltage pulse with 0 mM cAMP (filled circles, n = 36) and 1 mM cAMP (open circles, n = 15) in the intracellular solution. Sigmoidal fits (continues magenta lines), yielding midpoints of *I*_h_ activation (V_½_, arrows). The steady-state activation curves produced by the Hodgkin-Huxley models (dotted magenta line) are superimposed. *Inset on top*: Illustration of the whole-cell recording configuration with 0 and 1 mM cAMP in the intracellular solution. (C) Activation curve obtained with perforated-patch recordings show a shift in the *I*_h_ activation curve by 4.8 ± 1.2 mV compared to recordings from the same cell after rupture of the perforated membrane patch (n = 10). *Inset on top*: Illustration of the whole-cell recording configuration with 0 mM cAMP in the intracellular solution and in the perforated patch configuration, where the intracellular cAMP concentration is unperturbed. (D) Shift in *I*_h_ V_½_ versus the corresponding cAMP concentration (mean ± SEM). Fitting the data with a Hill equation revealed an EC_50_ of 40.4 µM. Superposition of the 68% confidence band of the fit (straight line and light magenta area) with the average voltage shift observed in perforated patch recordings (4.8 ± 1.2 mV, n = 10, dotted black line and grey area) results in an estimated cAMP-concentration in the non-perturbed presynaptic boutons of 12.6 µM with a range of 1.8 to 60.7 µM cAMP (dotted line and light blue area).

To analyze the cAMP concentration-dependence of *I*_h_, we added different concentrations of cAMP (30 µM to 10 mM) to the intracellular patch solution. With 1 mM cAMP V_½_ shifted by 17 mV to on average –86.6 ± 1.2 mV (n = 16 cMFBs; P_T-Test_ < 10^−10^; Fig. 8B). The resulting average shifts of V_½_ revealed an EC_50_ of 40.4 µM intracellular cAMP (Fig. 8D). In order to estimate the endogenous presynaptic cAMP concentration, we performed presynaptic perforated-patch recordings on cMFBs. Under perforated patch conditions, V_½_ of *I*_h_ was –96.4 ± 1.2 mV (n = 10), significantly more depolarized compared to the corresponding whole-cell recordings after rupturing of the perforated patch (–101.3 ± 1.0 mV; n = 15; P_T-Test_ = 0.0076; Fig. 8C; see methods for comparison with addition control groups). This voltage shift indicates an endogenous cAMP concentration of 12.6 µM in cMFBs, with a 68%-confidence interval of 1.8 to 60.7 µM cAMP (Fig. 8D). These data reveal a high endogenous resting cAMP concentration.

### Hodgkin-Huxley model describing HCN2 channel gating

For our ultimate aim, to obtain a mechanistic and quantitative understanding of axonal HCN2 function in cerebellar mossy-fiber axons, we developed a computational Hodgkin-Huxley (HH) model. The model was constrained to the experimentally recorded *I*_h_ kinetics derived from the activation and deactivation time constants of *I*_h_ (Fig. 9A) measured at potentials between –70 and –150 mV. The activation curve (cf. Fig. 9B), as well as the averaged time constants for both activation (n = 20) and deactivation (n = 15; Fig. 9B) were well described by a HH-model with one activation gate. In addition, we generated an alternative HH-model to describe the HCN2 current in the presence of 1 mM intracellular cAMP (for a more detailed implementation of the cAMP-dependence of HCN2 gating see Hummert et al., 2018). Furthermore, we estimated the reversal potential of *I*_h_ with short voltage ramps as described previously (Cuttle et al., 2001) and found a value of –23.4 ± 1.4 mV (n = 7; Fig. 9C), similar to previous estimates (Aponte et al., 2006; Cuttle et al., 2001). These data provide a quantitative description of axonal *I*_h_ at cMFBs.

**Figure 9.**
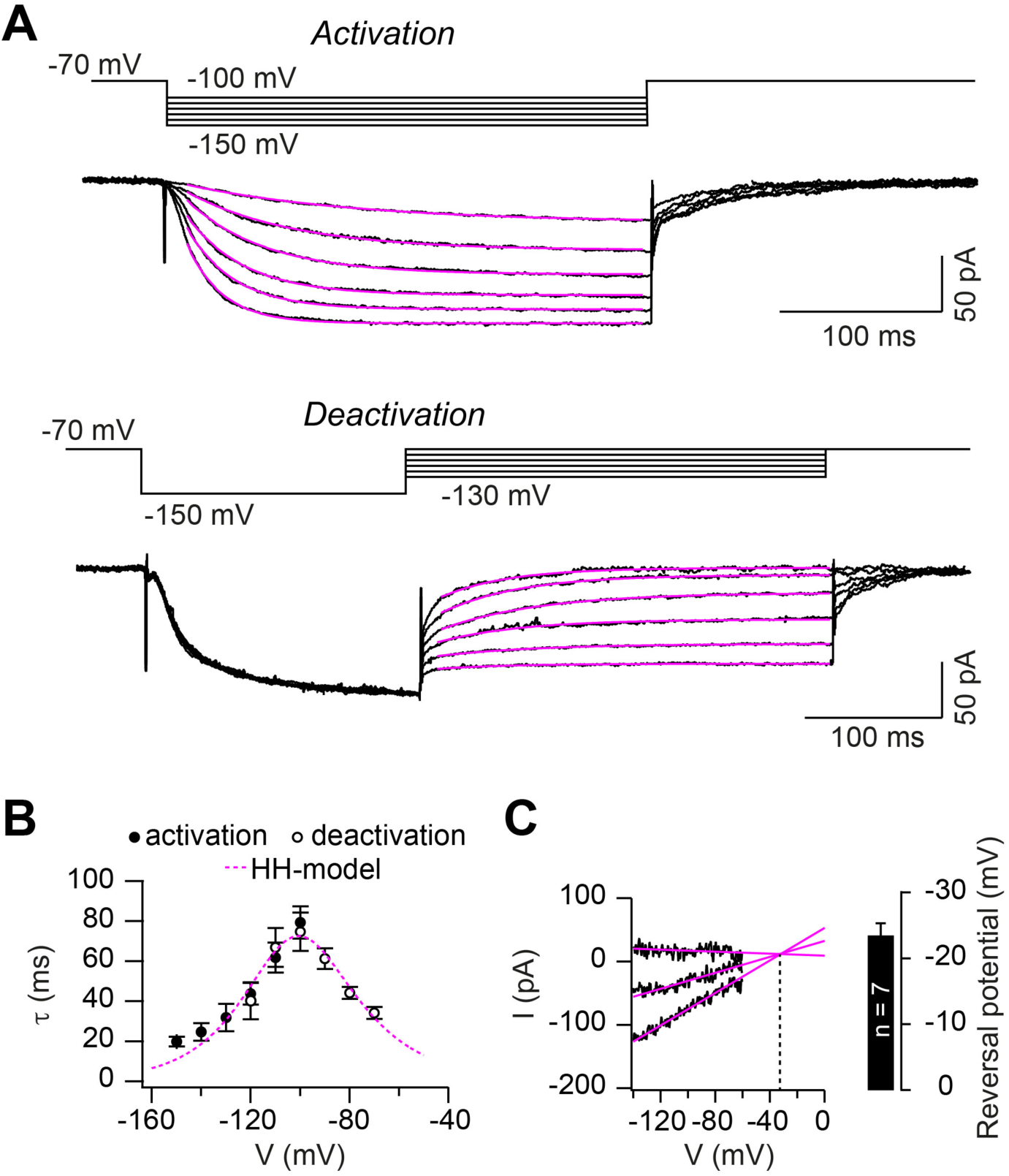
Hodgkin-Huxley model describing HCN2 channel gating. (A) Example of ZD7288 sensitive currents (black) elicited by the illustrated activation (*top*) and deactivation voltage protocols (*bottom*) superimposed with a mono-exponential fits (magenta). (B) Average time constants of activation (filled circles) and deactivation (open circles; mean ± SEM). The dotted blue line represents the prediction of *I*_h_ activation and deactivation time constant based on the Hodgkin-Huxley model. (C) Example of linear extrapolation (magenta lines) of leak subtracted currents evoked by fast (10 ms) voltage ramps generated from a range of holding potentials that extended across the activation range of *I*_h_. The reversal potential was found to be –36 mV in this example. *Inset:* average reversal potential of 7 independent experiments.

### Mechanism of conduction velocity-control and metabolic costs of HCN channels

What are the mechanisms by which axonal HCN2 channels control conduction velocity? In principle, the depolarization caused by HCN2 channels will bring the resting membrane potential closer to the threshold of voltage-gated Na^+^ channel activation, which could accelerate the initiation of the action potential (see discussion). Alternatively, the increased membrane conductance caused by HCN2 channels will decrease the membrane time constant, which could accelerate the voltage responses, as has been shown, e.g., for dendritic signals in auditory pathways (Golding and Oertel, 2012; Mathews et al., 2010). To distinguish between these two possibilities, we generated a conductance-based NEURON model consisting of cylindrical compartments representing cMFBs connected by myelinated axons (Fig. 10A; Ritzau-Jost et al., 2014). The model contained voltage-dependent axonal Na^+^ and K^+^ channels, passive Na^+^ and K^+^ leak channels, and the established HH model of *I*_h_ (cf. Fig. 9). After adjustments of the peak conductance densities the model captured the current clamp responses to –10 pA current injections (Fig. 10B), the resting membrane potential (Fig. 10C) as well as the apparent input resistance (Fig. 10D). Removing the HH model of *I*_h_ or replacing it with the 1-mM-cAMP-HH-model of *I*_h_, reproduced the corresponding voltage responses, the shift in the resting membrane potential, and the change in the apparent input resistance obtained in the presence of ZD7288 or 1 mM intracellular cAMP (Fig. 10B-D). Interestingly, the models predicted a decrease of the conduction velocity when the control HH model was removed and, conversely, an increase with the 1-mM-cAMP-HH model (Fig. 10E), to a similar extent as experimentally measured with ZD7288 and 8-Br-cAMP (cf. Fig. 1). These findings support our conclusion that HCN2 channel modulation suffices to bi-directionally tune conduction velocity.

**Figure 10.**
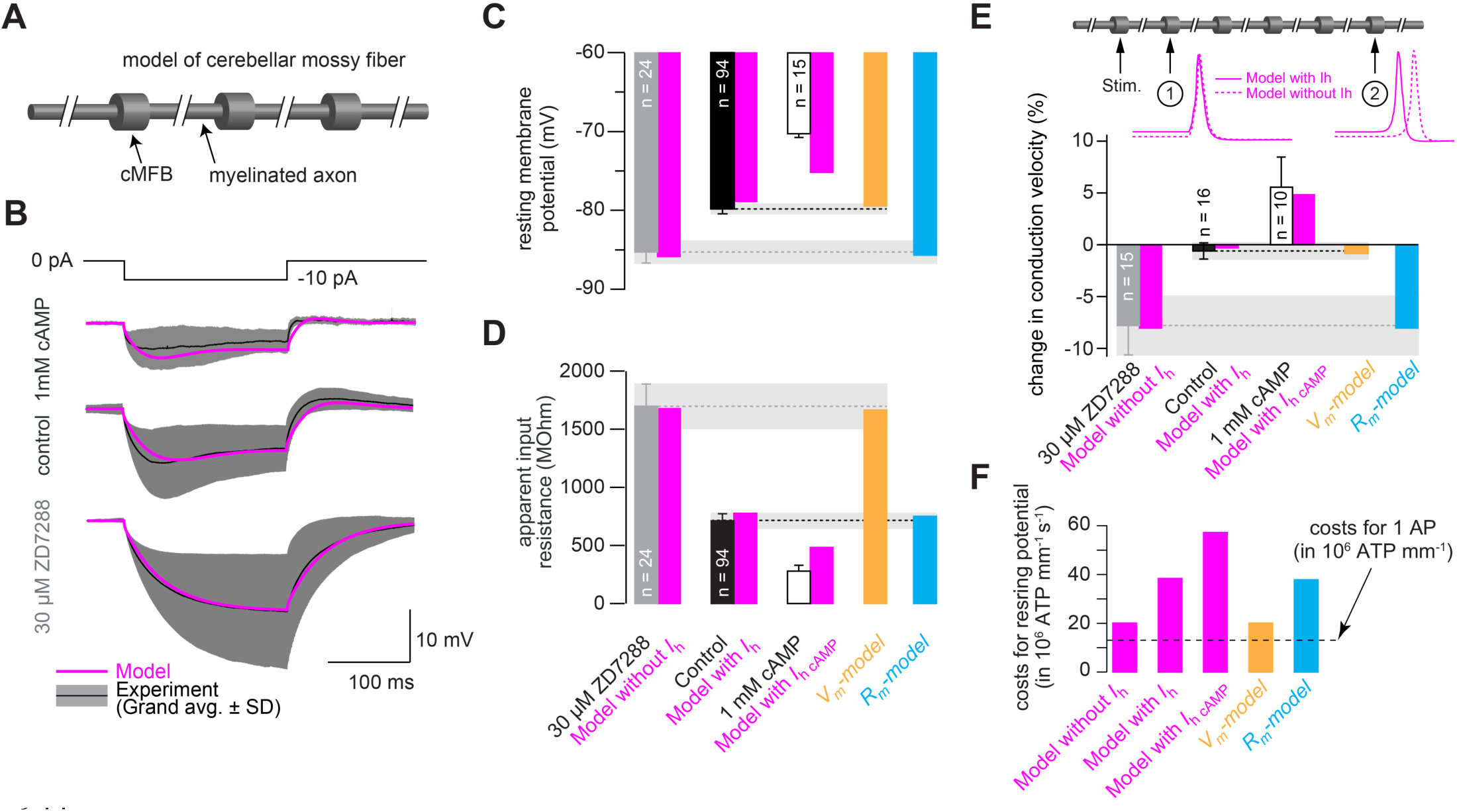
Mechanism of conduction velocity-control and metabolic costs of HCN channels. (A) Illustration of cerebellar mossy fiber model consisting of 15 connected cylindrical compartments representing cMFBs and the myelinated axon. (B) Grand average voltage response (black) and standard deviation (grey area) of cMFBs in response to a –10 pA hyperpolarizing current pulse with 1mM cAMP included in the patch pipette (top), under control conditions (middle) or treated with ZD7288 (bottom), superimposed with the predicted voltage response from the model (magenta). (C) Average resting membrane potential of cMFBs measured under control conditions (black), with *I*_h_ blocked by ZD7288 (gray), or enhanced by 1 mM intracellular cAMP (open bar; data from Fig. 6B) and compared to the predictions from the corresponding models (magenta). Furthermore, the resting membrane potential of two models is shown that simulate only the membrane depolarization (*V_m_-model*; light brown) or only the decreased membrane resistance (*R_m_-model*; blue) caused by open HCN channels. (D) Same comparison between measured values and predictions from the models for the apparent input resistance of cMFBs. (E) Measured changes of conduction velocity in mossy fibers compared to the predictions by the different models (as in C and D). *Inset top:* Illustration of the model of a mossy fiber with the stimulation positions and the action potentials at two different positions with (magenta line) and without (dashed magenta line) the HH model of HCN channels. (F) The calculated metabolic costs for maintaining the resting membrane potential are shown for each model as the number of required ATP molecules per mm of mossy fiber axon and per s. The metabolic cost for the firing of a single action potential (AP) is indicated by the dashed line as the number of required ATP molecules per mm of mossy fiber axon (this number was very similar for all models).

Next, we generated two additional models, in which either only the depolarizing effect of HCN2 channels (*V_m_-model*) or the decreased input resistance, i.e. the decreased membrane resistance (*R_m_-model*) was implemented (by modifying the K^+^ or the *I*_h_ reversal potential, respectively, see methods). The results showed that the *V_m_-model* but not the *R_m_-model* caused an increase in conduction velocity, indicating that the depolarizing effect of axonal HCN2 channels determines conduction velocity (Fig. 10E). Interestingly, increasing the resting membrane potential from –90 to –65, decreased the availability of voltage dependent Na^+^ (Na_V_) channels but increased the conduction velocity (Fig. S2). Only at resting membrane potentials above –65 mV the conduction velocity decreased in our model. Together, these data indicate that the depolarization mediated by HCN2 channels accelerates the conduction velocity by bringing the membrane potential closer to the firing threshold.

The non-inactivating nature of HCN channels and the accompanying shunt at the resting membrane potentials suggest that *I*_h_ is metabolically expensive. Therefore, we calculated the Na^+^ influx in each model and converted it into the required ATP consumption to restore the Na^+^ gradient (Hallermann et al., 2012). Computational modeling showed that it is ∼100% more expensive to maintain the resting membrane potential with *I*_h_ than without or by depolarization alone (*V_m_-model*; Fig. 10F). Furthermore, the metabolic costs to maintain the resting membrane potential with *I*_h_ for one second was ∼3-fold higher than the costs to generate one action potential (Fig. 10F). Assuming an average frequency of cerebellar mossy fibers of 4 Hz *in vivo* (Chadderton et al.; Rancz et al., 2007), the HCN2 channels increased the required energy of cerebellar mossy fibers by ∼30%. With increasing firing frequency, the metabolic costs of action potential firing will become dominating compared with the HCN2-mediated costs for resting membrane potentials (e.g., ∼3% at 40 Hz). These data indicate that HCN2 channels are a major contribution for the energetic demands of axons.

## Discussion

Here, we demonstrate that the presence of HCN channels in axons accelerates conduction velocity in various types of central axons and that neuromodulators change axonal conduction velocity. By combining advanced electrophysiological, electron-microscopic, and computational techniques, we reveal the mechanism and the metabolic costs of the dynamic control of axonal conduction velocity by HCN channels in the vertebrate central nervous system.

### Dynamic control of conduction velocity

We describe both an increase and decrease of the baseline axonal conduction velocity in the range of ∼5% mediated by HCN channels (Figs. 1 – 3). Furthermore, HCN channel accelerate action potential propagation by ∼25% and increase the maximal failure-free firing frequency by a factor of 2. Although the changes in baseline conduction velocity are relatively small, considering the long distances that axons traverse in the brain, HCN channels can be expected to change the arrival time of the action potential by for example 0.5 ms in the case of cerebellar parallel fibers (assuming 3 mm length and 0.3 m/s velocity; see also Swadlow and Waxman, 2012). Such temporal delays will influence information processing in the central nervous system, because spike-timing dependent plasticity (Caporale and Dan, 2008), coincidence detection (Softky, 1994), and neuronal rhythms of cell ensembles (Buzsáki et al., 2013) precisely tune the arrival times of action potentials. There are several examples for specific tuning of conduction velocity in the sub-millisecond domain: the diameter and the degree of myelination of cerebellar climbing fibers (Sugihara et al., 1993; Lang and Rosenbluth, 2003; but see Baker and Edgley, 2006), the degree of myelination of thalamocortical axons (Salami et al., 2003), and the internode distance of auditory axons (Ford et al., 2015) are tuned to exactly offset different arrival times of action potentials with a temporal precision of ∼100 µs.

The cerebellum is involved in the accurate control of muscle contraction with a temporal precision of 1-100 ms (Hore et al., 1991). Submillisecond correlations in spike timing occuring between neighboring Purkinje cells have been noted previously (reviewed in Isope et al., 2002; Person and Raman, 2012). Furthermore, submillisecond precision of the mossy/parallel fiber input are critical for information processing in the cerebellar circuits (Braitenberg et al., 1997; Heck et al., 2001). Together, the here-described changes in action potential conduction velocity in mossy and parallel fibers (Figs. 1–3) may thus play an important role in cerebellar computation. Furthermore, the observed impairment in high-frequency firing without HCN channels (Fig. 5) is expected to negatively impact such functions (Delvendahl and Hallermann, 2016).

Our findings that the cAMP-HCN pathway and neuromodulators can finely tune conduction velocity in the vertebrate central nervous system adds to the emerging idea that axons directly contribute to computation in neuronal circuits. Indeed, the view of the axon as a simple and cable-like compartment in which conduction velocity is static has substantially changed over the recent years in favor for a model that allows flexibility and complex forms of axonal computation (Debanne et al., 2011). Recent findings showed that axon diameters change during the time scales of LTP induction (Chéreau et al., 2017) and changes in myelination in the motor cortex were resolved during learning of complex motor skills (McKenzie et al., 2014; for environmental effects on myelination see also Forbes and Gallo, 2017). One caveat of our study, is the rather high concentrations of the used neuromodulators and the lack of *in vivo* evidence for neuromodulation of conduction velocity. However, our data demonstrate that under certain conditions an active control of conduction velocity could occur in the vertebrate CNS via the cAMP-HCN pathway.

### Mechanism and metabolic costs of HCN channel mediated control of conduction velocity

Our analysis revealed that the control of conduction velocity is solely mediated by changes in resting membrane potential. Isolated changes of membrane conductance and thus of the membrane time and length constant had no effect on conduction velocity (Fig. 10E). The speeding of conduction upon depolarization is consistent with a previously observed correlation between conduction velocity and the depolarization from the resting potential required to reach the firing threshold in motoneurons (Carp et al., 2003). On the other hand, Na^+^ channels have a steep steady-state inactivation and are partially inactivated at the resting membrane potential in axons (Battefeld et al., 2014; Engel and Jonas, 2005; Rama et al., 2015). Depolarization could thus be expected to further inactivate Na^+^ channels and decrease conduction velocity. However, our modelling results showed that increasing the membrane potential from –90 to –60 mV increased the conduction velocity despite significantly decreasing Na^+^ channels availability (Fig. S2). Interestingly, these findings are in agreement with the nonlinear cable theory predicting that the difference between the resting membrane potential and the firing threshold is a critical parameter for action potential conduction velocity (see, e.g., Fig. 12.25 in Jack et al., 1983, for increasing velocity with increasing safety factor, i.e. decreasing excitation threshold *V*_B_). Intuitively, the HCN channel mediated acceleration of conduction velocity can be understood as follows; in a more depolarized axon, Na_V_ mediated current influx in one axonal location will depolarize neighboring locations faster above the threshold. In our model, this effect outweighs the disadvantage of the increased steady-state inactivation of Na^+^ channels up to a membrane potential of about –65 mV and a Na_V_ availability of 50%. The exact values, above which Na_V_ availability limits conduction velocity, critically depend on assumptions of the model, such as the voltage-dependence of inactivation and the density of Na_V_ channel. Interestingly, Ca^2+^ entering through axonal voltage-gated Ca^2+^ channels (Bender et al., 2010) could interact with the cAMP pathway by activating or inhibiting different subtypes of adenylyl cyclase and phosphodiesterase (Bruce et al., 2003). Thereby, HCN could also serve a homeostatic role, by bringing the resting membrane potential closer to threshold and offsetting the inactivation of Na_V_ channels under conditions of high-frequency action potential firing.

Our findings indicate that the evolutionary design of HCN channels as a continuously open shunt for Na^+^ influx causes significant metabolic costs. The high costs might appear surprising, because a metabolically cheaper way to depolarize the membrane would be the expression of less Na^+^-K^+^-ATPases resulting in a depolarized K^+^ reversal potential (cf. *V_m_-model* in Fig. 10). However, as discussed in the following paragraph, our finding that conduction velocity can be rapidly regulated via the cAMP-HCN pathway might provide a justification for the metabolic costs of axonal HCN channels.

### Modulation of conduction velocity via the intracellular cAMP concentration

Using direct whole-cell recordings and immunogold EM from *en passant* boutons in cerebellar axons, we identified near exclusive HCN2 isoforms expression and a half-maximal shift of the activation of HCN2 channels at a cAMP concertation of 40 µM (Fig. 7D). Furthermore, our perforated patch-recordings from axonal compartments provide, to our knowledge, the first direct estimate of endogenous cAMP concentration in vertebrate central axons of 13 µM (Fig. 7D). This is higher compared to previous estimates of 50 nM in Aplysia sensory neurons (Bacskai et al., 1993; but see Greenberg et al., 1987) and 1 µM in cardiomyocytes (Börner et al., 2011). A recently reported low cAMP-sensitivity of protein kinase A (Koschinski and Zaccolo, 2017), a prototypical cAMP-regulated protein, also argues for high intracellular cAMP concentrations. On the other hand, our data do not rule out that such high cAMP concentration occur only in spatially restricted domains. The possibility for local cAMP signaling-compartments was recently observed in *Drosophila* axons (Maiellaro et al., 2016).

A high endogenous cAMP concentration and expression of the HCN2 isoform facilitates neuromodulators to bidirectionally and dynamically control conduction velocity. Only norepinephrine increased the conduction velocity in cerebellar parallel fibers whereas the other neuromodulators reduced the velocity (Fig. 2), consistent with the expression of both G_i_- and G_s_-coupled receptors, respectively. Indeed, G_s_-coupled receptors for serotonin, dopamine, and adenosine are expressed in the molecular layer of the cerebellum (see e.g. Geurts et al., 2002; Schweighofer et al., 2004). Interestingly, adenosine, which decreased the conduction velocity (Fig. 2), has been shown to be an endogenous sleep factor (Basheer et al., 2004; Porkka-Heiskanen et al., 1997). Moreover, serotonin, dopamine, and norepinephrine play important regulatory functions during sleep in, e.g., the cerebellum (Canto et al., 2017). Therefore, it is tempting to speculate that the cAMP-HCN pathway allows not only the increase in conduction velocity during arousal but also the decrease in velocity and saving of metabolic costs during periods of rest or sleep. The cAMP-HCN pathway in axons could thus contribute to the reduced energy consumption of the brain during sleep (Boyle et al., 1994; Townsend et al., 1973). It should be noted that the observed modulation of conduction velocity by neurotransmitters (Fig. 2) is consistent with a modulation via the cAMP-HCN pathway but other mechanisms, such as direct influences on voltage-dependent Na^+^ (Yin et al., 2017), K^+^ (Yang et al., 2013), and Ca^2+^ channels (Burke et al., 2018) could contribute to the modulation of conduction velocity.

### Clinical relevance of axonal HCN channels

The function of HCN channels has been studied in human peripheral nerves using non-invasive threshold tracking techniques (Howells et al., 2016; Howells et al., 2012; Lorenz and Jones, 2014). Significant alterations of HCN channel expression and/or function have been described in pathologies such as stroke (Jankelowitz et al., 2007), porphyria (Lin et al., 2008), diabetic neuropathy (Horn et al., 1996), neuropathic pain (Chaplan et al., 2003), and inflammation (Momin and McNaughton, 2009) as well as a vertebrate model of demyelination (Fledrich et al., 2014). In some of these cases, the alterations are consistent with an activity-dependent modulation of HCN channels (Jankelowitz et al., 2007). Furthermore, HCN channel seem to be causally related to pain symptoms (Chaplan et al., 2003; Momin and McNaughton, 2009) and therapeutic blockade of HCN channels are also considered (Wickenden et al., 2009). Based on our findings, HCN could also play a compensatory role in some diseases to restore conduction velocity.

## Methods

### Cerebellar slice preparation

Cerebellar slices were prepared from P21-P46 C57BL/6 mice of either sex as reported previously (Delvendahl et al., 2015; Ritzau-Jost et al., 2014). In short, after anesthetization with isoflurane, mice were killed by rapid decapitation; the cerebellar vermis was quickly removed and placed in a slicing chamber filled with ice-cold extracellular solution (ACSF) containing (in mM): NaCl 125, KCl 2.5, NaHCO_3_ 26, NaH_2_PO_4_ 1.25, glucose 20, CaCl_2_ 2, MgCl_2_ 1 (pH adjusted to 7.3–7.4 with HCl). Parasagittal or horizontal slices were cut from the vermis of the cerebellum using a microtome with a vibrating blade (VT1200, Leica Biosystems, Nussloch, Germany), incubated at 35°C for approximately 30 minutes and subsequently stored at room temperature until use. For electrophysiological recordings, a slice was transferred into the recording chamber mounted on the stage of an upright Nikon microscope. The recording chamber was perfused with ACSF and the temperature in the center of the recording chamber was set to 35°C using a TC-324B perfusion heat controller (Warner Instruments, Hamden CT, USA).

### Measuring conduction velocity in cerebellar parallel and mossy fibers

Compound action potentials were evoked by electrical stimulation using a bipolar platinum/iridium electrode (from Microprobes for Life Science, Gaithersburg MD, USA) placed either in the white matter or in the molecular layer (Fig. 1) of the cerebellum. For the extracellular recording of compound action potentials, two pipettes were filled with a 1M NaCl solution (tip resistance of 1–3 MΩ) and placed within the respective fiber bundle, and the voltage was measured in current clamp mode with an EPC10 amplifier (CC gain 10x). Compound action potentials were evoked at 0.5 Hz in parallel fibers and 1 Hz in the white matter. All recordings were performed in the presence of 10 µM NBQX to block synaptic potentials. The conduction velocity of PFs was measured at 35°C. Due to the higher conduction velocity in myelinated mossy fibers, action potentials evoked by white matter stimulation had to be recorded at room temperature to allow separation of the compound action potential from the stimulation artifact. To calculate the conduction velocity, we determined the delays of the peaks of the compound action potential component recorded with the proximal and distal electrode. Compound action potentials from mossy fibers were analyzed offline using the smoothing spline interpolation operation of Igor Pro to increase the signal to noise ratio. Control recordings were performed interleaved with application of different drugs. The conduction velocity experienced a small rundown over 20 minutes under control conditions (Fig. 1 – 3).

### Measuring conduction velocity in the optic nerve

Male wildtype mice of the C57BL6/N strain (P63 ± 4) were euthanized by decapitation. After the brain was exposed, the optic nerves (ON) were separated from the retina at the ocular cavity and both ONs were detached by cutting posterior to the optic chiasm. The preparation was gently placed into an interface brain/tissue slice (BTS) perfusion chamber (Harvard Apparatus) and continuously superfused with ACSF, bubbled with carbogen (95% O_2_, 5% CO_2_) at 36.5 °C during the experiment (Trevisiol et al., 2017). In case both nerves were used for experiments, the non-recorded ON was transferred in a different incubation chamber (Leica HI 1210) that provided similar incubation conditions to the recorded nerve while preventing exposure to ZD7288 and 8-Br-cAMP. The temperature was maintained constant using a feedback-driven temperature controller (model TC-10, NPI electronic) connected to a temperature probe (TS-100-S; NPI electronic) inserted in the BTS incubation chamber near the nerve. Each ON was detached from the optic chiasm and individually placed into the suction electrodes for stimulation/recording. The stimulation’s direction of the ON was maintained constant (orthodromic) throughout the experiments by inserting the proximal (retinal) end of the nerve into the stimulation electrode as illustrated in Fig. 1I. The stimulating electrode was connected to a battery (Stimulus Isolator A385; WPI) that delivered a supramaximal stimulus to the nerve. The voltage was pre-amplified 500 times and fed to the AD ports of the EPC9 or acquired directly via the EPC9 headstage (HEKA Elektronik, Lambrecht/Pfalz). The reference channel was obtained from an ACSF-filled glass capillary next to the recording suction electrode, in contact with the bathing ACSF. Initial equilibration of the ONs was performed at 0.1 Hz stimulation, until the recorded compound action potentials showed a steady shape (typically around 45-60 min from preparation). 5 nerves from 4 animals and 4 nerves from 4 animals were used for ZD7288 and 8-Br-cAMP treatment, respectively. Compound action potentials were analyzed as described above using the smoothing spline interpolation operation of Igor Pro to increase the signal to noise ratio.

### Recordings from cMFBs

cMFBs were visualized as previously described (Delvendahl et al., 2015; Ritzau-Jost et al., 2014) with infrared differential interference contrast (DIC) optics using a FN-1 microscope from Nikon with a 100x objective (NA 1.1) or infrared oblique illumination optics using a Femto-2D two-photon microscope (Femtonics, Budapest) with a 60x Olympus (NA 1.0) objective. The passive properties of the cMFB were determined as previously described (Hallermann et al., 2003) and revealed similar values for a two-compartment model (data not shown) as previously described for cMFBs (Ritzau-Jost et al., 2014), indicating that we indeed recorded from cMFBs. Furthermore, the access resistance was on average 16.9 ± 0.9 MΩ (n = 53 cMFBs), indicating optimal voltage clamp conditions.

To elicit traveling action potentials by axonal stimulation with a second pipette (Fig. 5A), whole-cell recordings from cMFBs were performed with 50 µM green fluorescent dye Atto488 in the intracellular solution to visualize single mossy fiber axons. The additional stimulation pipettes filled with ACSF and 50 µM of the red-fluorescent dye Atto594 had the same opening diameter as patch pipettes and were positioned close to the axon and approximately 100 µm apart from the patched terminal. Stimulation pulses with durations of 100 µs were delivered by a voltage-stimulator (ISO-Pulser ISOP1, AD-Elektronik, Buchenbach, Germany). The stimulation intensity (1-30 V) was adjusted to ensure failure-free initiation of action potentials at 1 Hz (∼1.5 time the firing threshold). High-frequency trains of action potentials were evoked at 100, 200 333, 500, 750, 1000, 1111 and 1666 Hz. Amplitudes were measured from peak to baseline. The duration was determined at half-maximal amplitude and is referred to as half-width. Action potentials were treated as failures if the peak did not exceed –40 mV.

Recordings were performed with an EPC10/2 patch-clamp amplifier, operated by the corresponding software PatchMaster (HEKA Elektronik), running on a personal computer. Recording electrodes were pulled from borosilicate glass capillaries (inner diameter 1.16 mm, outer diameter 2 mm) by a microelectrode puller (DMZ-Universal Puller, Zeitz Instruments, Augsburg). Pipettes used for patch-clamp recordings had open-tip resistances of 5–12 MΩ. The intracellular presynaptic patch pipette contained (in mM): K-gluconate 150, MgATP 3, NaGTP 0.3, NaCl 10, HEPES 10 and EGTA 0.05. The apparent input resistance of cMFBs was estimated by linear regression of the steady-state voltage in response to 300 ms hyperpolarizing current pulses of increasing amplitude (–5 to –20 pA), while the apparent membrane time constant was determined by fitting the voltage response to a −10 pA hyperpolarizing pulse with a mono-exponential function.

*I*_h_ activation curves determined from analysis of normalized tail current were fitted with a Boltzmann function:

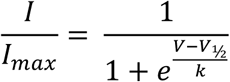

where V is the holding potential, *V_½_* is the voltage of half-maximal activation and *k* the slope factor. The reversal potential of *I*_h_ was calculated from leak-subtracted currents evoked by 10 ms long voltage ramps extending across the activation range of *I*_h_ (Cuttle et al., 2001). Three I-V relationships recorded at holding potentials of –80, –110 and – 140 mV were linearly extrapolated and the reversal potential was measured from the point of intersection of the three linear fits.

### Perforated patch recordings

For perforated-patch recordings from cMFBs, a nystatin stock solution was prepared by dissolving the pore-forming antimycotic in DMSO (25 mg/ml). Immediately before the experiments, the nystatin-stock was added to the intracellular solution at a final concentration of 50 µg/ml. In order to monitor the integrity of the perforated membrane patch, the green-fluorescent dye Atto 488 (from Atto-Tec, Siegen, Germany) was added at a concentration of 50 µM. Since nystatin is known to impair the formation of the GΩ seal, the initial ∼500 µm of the pipette tip was filled with a perforating agent-free internal solution before back-filling the pipette shaft with the perforating agent-containing solution. After establishing a GΩ seal, the holding potential was set to –70 mV and the access resistance (R_a_) was continuously monitored by applying 10 ms long depolarizing pulses to –60 mV at 1 Hz. Recording the voltage-dependent activation of *I*_h_ was started after R_a_ dropped below 150 MΩ. Because the perforated membrane patch ruptured spontaneously with R_a_ < 50 MΩ, the access resistance was not comparable to standard whole-cell recordings. To exclude the possibility that the right-shift of the *I*_h_ activation curve in the perforated configuration (Fig. 8C) was caused by the comparatively higher R_a_, the voltage-dependent activation of *I*_h_ was measured under normal whole-cell patch-clamp conditions, using pipettes with small openings resulting in high access resistances (R_a_ = 119 ± 12 MΩ). However, in these recordings, the midpoint of *I*_h_ activation (–105.5 ± 1.4 mV; n = 8) had a tendency to be left-shifted compared with regular whole-cell recordings with standard patch pipettes (R_a_ ≈ 30-60 MΩ; V_½_ = –103.3 ± 0.8 mV; n = 36; P_T-Test_ = 0.13). The left-shift of the *I*_h_ activation curve measured with high access resistances indicates that the right-shift measured with perforated patch recordings might be underestimated due to the higher R_a_, which would result in an even higher estimate of the endogenous cAMP concentration (Fig. 7D).

### Analysis of ZD sensitivity of Na^+^ currents

Sodium currents (Fig. S1) were isolated using a modified ACSF containing (in mM): NaCl 105, KCl 2.5, NaHCO_3_ 25, NaH_2_PO_4_ 1.25, glucose 20, CaCl_2_ 2, MgCl_2_ 1, TEA 20, 4-AP 5 and CdCl_2_ 0.2. To avoid underestimating the true size of the presynaptic Na^+^ currents due to the voltage-drop through the access resistance, we blocked a portion of the Na^+^ current with 30 nM TTX. Na^+^ currents were elicited from a holding potential of –80 mV by a 3 ms long depolarization to 0 mV. Peak amplitudes and half-durations of Na^+^ currents were measured from leak-subtracted traces.

### Immunoelectron microscopy

Preembedding immunogold labeling was performed as described (Notomi and Shigemoto, 2004). Briefly, adult C57Bl/6 mice were anesthetized with sodium pentobarbital (50 mg/kg, i.p.) and perfused transcardially with a fixative containing 4% formaldehyde, 0.05% glutaraldehyde and 15% of a saturated picric acid in 0.1 M phosphate buffer (PB, pH 7.4). Parasagittal sections through the cerebellum were cut at 50 µm, cryoprotected with 30% sucrose, flash frozen on liquid nitrogen and rapidly thawed. Sections were blocked in 10% normal goat serum and 2% bovine serum albumin (BSA) in Tris-buffered saline (TBS) for 2 h at room temperature, incubated in TBS containing 2% BSA and either guinea pig anti-HCN1 or anti-HCN2 antibody (1 µg/ml, Notomi and Shigemoto, 2004) for 48 h at 4°C, and finally reacted with nanogold-conjugated secondary antibody (Nanoprobes, 1:100) for 24 h at 4°C. Nanogold particles were amplified with HQ Silver Enhancement kit (Nanoprobes) for 8 min. Sections were then treated in 0.5% osmium tetroxide in PB for 40 min, 1% aqueous uranyl acetate for 30 min at room temperature, dehydrated, and flat embedded in Durcopan resin (Sigma-Aldrich). Ultrathin sections were cut at 70 nm and observed by a transmission electron microscope (Tecnai 12, FEI, Oregon). Sequential images were recorded from the granule cell layer within a few microns from the surface of ultrathin sections at X26,500 using a CCD camera (VELETA, Olympus). For the reconstruction of a half mossy fiber bouton, 36 serial ultrathin sections were used. Sequential images were aligned and stacked using TrakEM2 program (Cardona et al., 2012). For the measurement of density of immunogold particles for HCN2 on this reconstructed profile, 1260 immunogold particles were counted on the mossy fiber bouton membrane area (73.7 µm^2^), giving a density of 17.1 particles/µm^2^. Immunogold particles within 30 nm from the bouton membranes were included in the analysis based on the possible distance of the immunogold particle from the epitope (Matsubara et al., 1996). The density of non-specific labeling was estimated using nuclear membrane of a granule cell located adjacent to the reconstructed mossy fiber bouton. We found 40 immunogold particles on the nuclear membrane area of 60.5 µm^2^ giving a density of 0.66 particles/µm^2^, which was 3.9% of the HCN2 labeling density on the mossy fiber bouton.

### Hodgkin-Huxley model of axonal HCN channels

Because we did not intend to implement the cAMP dependence of HCN channels explicitly (Hummert et al., 2018), we created two separate models for 0 and 1 mM intracellular cAMP, which were based on a previously described Hodgkin-Huxley model (Kole et al., 2006) with one activation gate and no inactivation (Hodgkin and Huxley, 1952). In short, the activation gate was described by

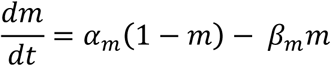

with

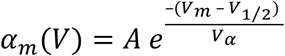

and

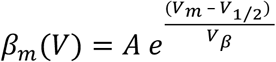

The four free parameters, *A*, *V_1/2_*, *V_α_*, and *V_β_* were determined by simultaneously fitting *α_m_* /(*α_m_* **+** *β_m_*) to the steady-state activation curve (see Fig. 7B) and 1/(*α_m_* **+** *β_m_*) to the voltage dependence of the time constant of *I*_h_ activation and deactivation (Fig. 9B). The sum of squared errors was minimized using the FindMinimum routine of Mathematica (version 10; Wolfram Research, Champaign, IL), with the time constants of activation and deactivation weighed with the inverse of the square of the maximum value in each of the three datasets (time constant of activation, time constant of activation, steady-state activation curve). The resulting parameters for 0 mM cAMP were *A* = 6.907 ms^−1^, *V_1/2_* = –102.1 mV, *V_α_* = 18.71 mV, and *V_β_* = 21.73 mV. To confirm that the global minimum was reached, the best-fit parameters were shown to be independent of the starting values within a plausible range. The 68% confidence interval was calculated as the square roots of the diagonals of the inverse of the Hessian matrix (Press et al., 2002) resulting in ±2.71 ms^-1^, ±16.5 mV, ±17.5 mV, and ±24.3 mV for *A*, *V_1/2_*, *V_α_*, and *V_β_*, respectively. We also generated a model for the corresponding data obtained with 1 mM cAMP in the intracellular solution (cf. Fig 5B), resulting in *A* = 7.570 ms^-1^, *V_1/2_* = – 87.31 mV, *V_α_* = 31.46 mV, and *V_β_* = 10.84 mV.

### NEURON model of cMFB

The model of the cMFB consisted of connected cylindrical compartments representing 15 boutons (length and a diameter 8 µm) and 15 myelinated axonal compartments (length 35 µm and a diameter 0.8 µm; cf. Palay and Chan-Palay, 1974; Fig. 10A). In addition, at one side of this chain a long cylinder was added presenting the axon in the white matter (length 150 µm and a diameter 1.2 µm). The specific membrane resistance was 0.9 µF/cm^2^ (Gentet et al., 2000) and the cytoplasmatic resistivity was 120 Ω/cm (Hallermann et al., 2003). The specific membrane resistance of the axonal compartments was reduced by a factor of 10 representing myelination.

The active membrane conductances were similar to Ritzau-Jost et al. (2014) and were adjusted to reproduce the action potential duration and maximal firing frequency as well as the data shown in Figs. 10B-D. Namely, an axonal Na^+^ channel (Schmidt-Hieber and Bischofberger, 2010) and K^+^ channel NMODL model (Hallermann et al., 2012) was added with a density of 2000 and 1000 pS/µm^2^ in the boutons and 0 and 0 pS/µm^2^ in the axonal compartments, respectively. The Na^+^ and K^+^ reversal potentials were 55 and –97 mV, respectively. To investigate ATP consumption, separate Na^+^ and K^+^ leak channel models were added, with a conductance of 0.0138 and 0.18 pS/µm^2^, respectively, in the bouton compartments. In the axonal compartments, both conductances were reduced by a factor of 10. The above described Hodgkin-Huxley model of axonal HCN channels for either 0 or 1 mM intracellular cAMP was added with a density of g_HCN_ = 0.3 and 0.03 pS/µm^2^ for the bouton and axonal compartments, respectively, to reproduce the data shown in Fig. 10B-D. To investigate ATP consumption, the conductance was separated in a Na^+^ and a K^+^ conductance according to g_HCN(Na)_ = (1 – ratio_K/Na_) g_HCN_ and g_HCN(K)_ = ratio_K/Na_ g_HCN_, where ratio_K/Na_ = (e_Na_ + e_HCN_)/(e_Na_ – e_K_), where e_Na_ and e_K_ are the Na^+^ and K^+^ reversal potential as described above and e_HCN_ is the reversal potential of *I*_h_ measured as –23.3 mV (cf. Fig. 9C). Assuming a single channel conductance of 1.7 pS for HCN2 channels (Thon et al.), this conductance corresponds to a density of 0.18 HCN channels/µm^2^, which is much lower than the estimate from preembedding immunogold labeling (22 particles/µm^2^; Fig. 8). However, the optimal density of the model critically depends on the geometry of the structure, which was not obtained from the recorded boutons. To obtain the required structural information including the fenestration of the cMFB (cf. Fig. 8) and the level of myelination, electron microscopic reconstructions of large volumes of the recorded cMFB and the entire axon would be needed. When we used a g_HCN_, as determined with preembedding immunogold labeling in our model, the model also predicted that *I*_h_ critically effects conduction velocity and that the depolarization is the main reason for the velocity to change. In general, these two conclusions of the model were very insensitive to the specific parameters of the model and were, e.g., also obtained with additional interleaved cylindrical compartments with high Na^+^ and K^+^ channel density representing nodes of Ranvier or with a long cylindrical compartment with homogenous channel densities representing an unmyelinated axon. This further supports our finding that HCN channels accelerate conduction velocity independent of the exact parameters of the axon and the degree of myelination (cf. Fig. 1).

Starting from the model that reproduced the control data, the following four additional models were generated: (1) To simulate ZD application, the HCN HH model was removed. (2) To simulate 8-br-cAMP application, parameters of the HCN HH model were exchanged with the parameters obtained from the experiments with 1 mM cAMP as described in the section above. (3) To simulate only the depolarization by HCN channels (*V_m_-model*), the HCN HH channel model was removed and the K^+^ reversal potential was increased from –97 mV to –90 mV. (4) To simulate only the increase in membrane conductance by HCN channels (*R_m_-model*), the reversal potential of the HCN HH model was decreased from –23.3 mV to –85.5 mV and the density was increased from 0.3 pS/µm^2^ to 1 pS/µm^2^.

All simulations were run with a simulation time interval (*dt*) of 0.2 ms, preceded by a simulation of 1 s with a *dt* of 5 ms to allow equilibration of all conductances. Conduction velocity was calculated from the peak of the action potentials in different boutons of the model. The apparent input resistance was calculated identical to the experimental recordings, i.e. from the voltage after 300 ms of a –10 pA current injection. Mathematica was used to execute the NEURON simulations and to visualize and analyze the automatically imported NEURON results.

### Statistics

Statistical analysis was performed using built-in functions of Igor Pro (Wavemetrics, Lake Oswego, OR). The suffix of the P values provided in the legends and the main test indicate the used statistical test. Results were considered significant with P < 0.05.

### Code

The NEURON and Mathematica scripts to reproduce the model results will be available at: https://github.com/HallermannLab/2018_eLife_HCN.

## Acknowledgement

We would like to thank Klaus Nave and the department of Neurogenetics at the Max Planck Institute of Experimental Medicine for scientific support. This work was supported by the German Research Foundation (HA 6386/4-1) to S.H.

---

**Supplemental Figure S1.**
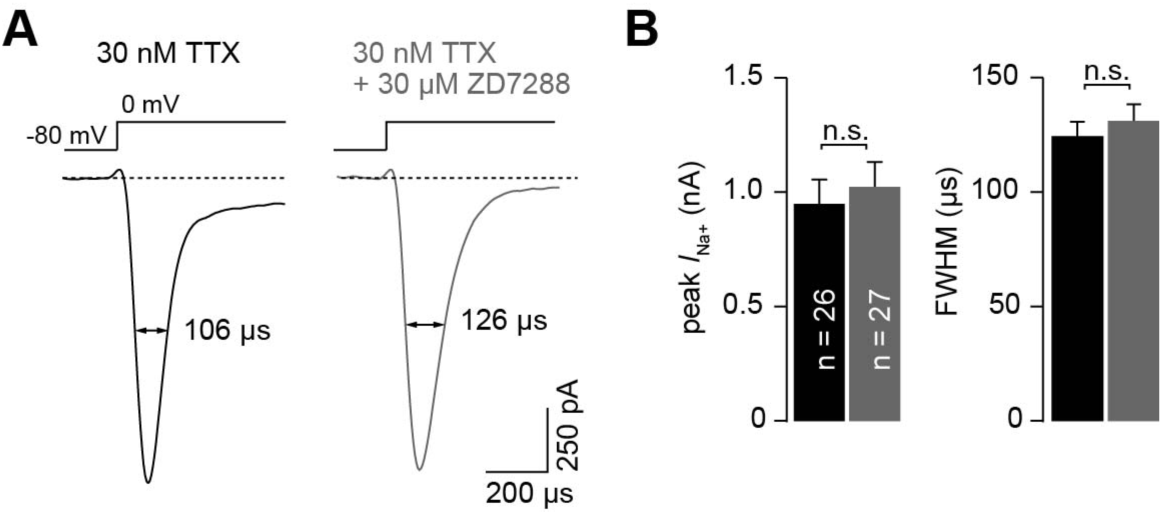
ZD7288 does not alter Na^+^ currents in cMFBs. (A) Example whole-cell Na^+^ currents measured under control conditions (30 nM TTX) and in the presence of additional 30 µM ZD7288 elicited by voltage steps from –80 mV to 0 mV. (B) With 30 nM TTX present in the extracellular solution (see methods), the average Na^+^ current amplitude was 947.9 ± 107.5 pA (n = 26). In the presence of 30 nM TTX and 30 µM ZD7288, the amplitude was not different compared with the currents recorded in 30 nM TTX only (1022.5 ± 109.0 pA; n = 27; P_T-Test_ = 0.63). The average half-duration was 124.3 ± 6.2 µs and 131.5 ± 7.2 µs (P_T-Test_ = 0.45) for control recordings and those performed in the presence of 30 µM ZD7288, respectively.

**Supplemental Figure S2.**
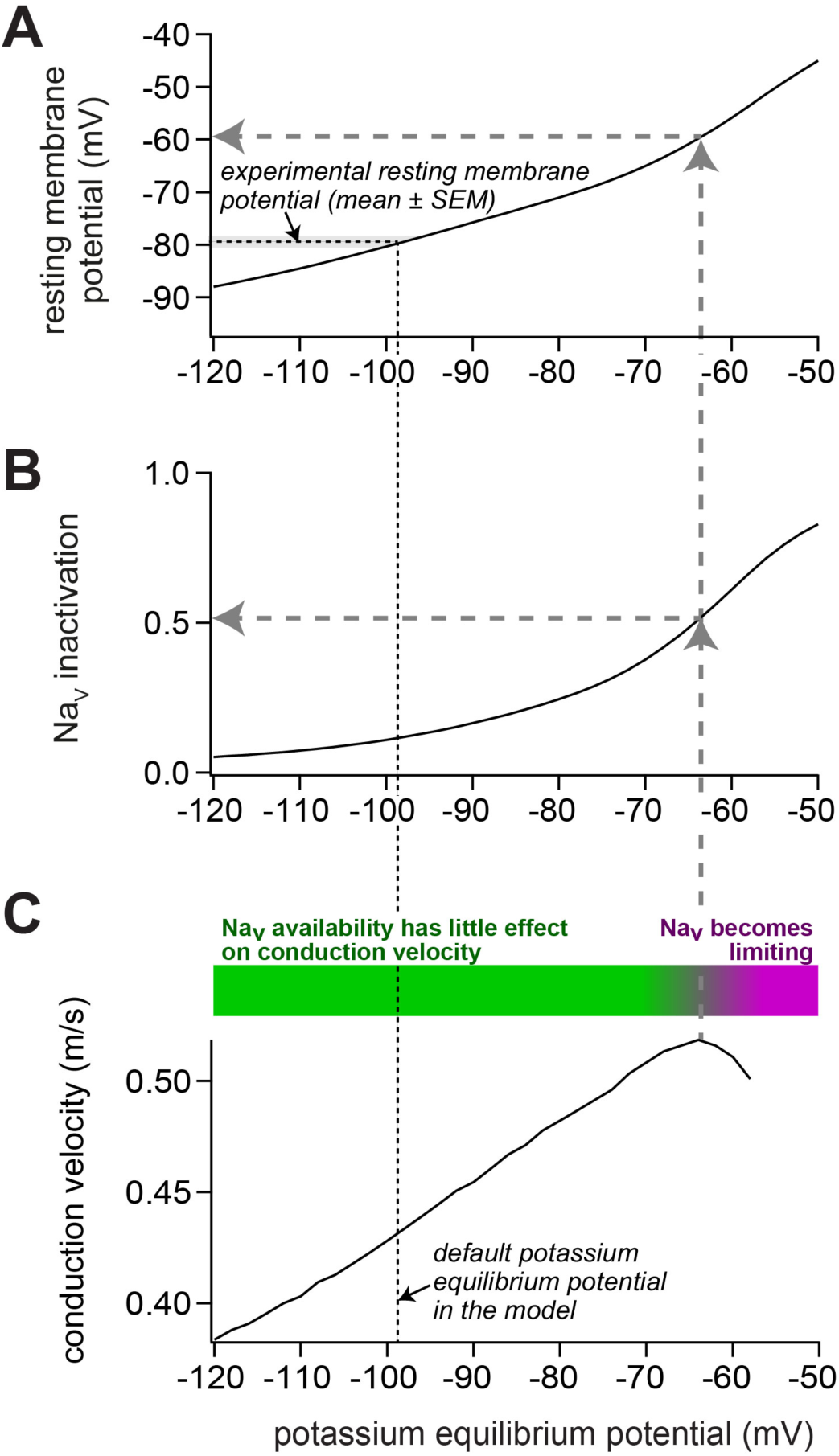
Impact of depolarization on Na_V_ availability and on conduction velocity in our model of a mossy fiber axon. (A) To change the resting membrane potential, the potassium equilibrium potential was varied between –120 and –50 mV. In the default model, the experimentally observed resting membrane potential (indicated by horizontal black dashed line and grey bar; mean ± SEM) was obtained with a potassium equilibrium potential of –98 mV (vertical black dashed line). (B) Upon depolarization, the inactivation of voltage-dependent sodium (Na_V_) channels increased. For the control model, the Na_V_ inactivation was 12%. For the model reproducing the experiments with ZD7288 and 1 mM intracellular cAMP, the Na_V_ inactivation was 6 and 17%, respectively. Because the steady-state Na_V_ availability depends mostly on the resting membrane potential, the relation between resting membrane potential and Na_V_ availability was identical for all three models (data not shown). (C) Upon depolarization, the conduction velocity increased up to a potassium equilibrium potential of about –65 mV (vertical grey dashed arrow), corresponding to an Na_V_ availability of about 50% (horizontal grey dashed arrow in panel B) and a resting membrane potential of about –60 mV (horizontal grey dashed arrow in panel A). Thus, despite increasing Na_V_ inactivation, the conduction velocity increased in this range (illustrated by green bar). With stronger depolarization, the conduction velocity declined, indicating that the Na_V_ availability becomes limiting for conduction velocity (magenta bar).

## Supplemental video 1 (video1.mp4). Reconstructed cMFB with labelled synapses and HCN2 channels

3D rendering of a part of a reconstructed cMFB (red) with identified synapses (blue) and HCN2 labeled with gold particles (yellow) based on immune-gold electron microscopic images.

